# Diversity, Phylogenetic Relationships, And Expression Profiles Of Invertase Inhibitor Genes In Sweetpotato

**DOI:** 10.1101/2022.08.31.505262

**Authors:** Samuel Acheampong, Heike Sederoff, Bode A. Olukolu, Aaron T. Asare, G. Craig Yencho

**Affiliations:** Department of Molecular Biology and Biotechnology, University of Cape Coast, Cape Coast, Ghana; Department of Plant and Microbial Biology, North Carolina State University, Raleigh, North Carolina, USA; Department of Entomology & Plant Pathology, The University of Tennessee, Knoxville, Tennessee, USA; Department of Horticultural Science, North Carolina State University, Raleigh, North Carolina, USA

**Keywords:** invertase inhibitor, sweetpotato, sequence diversity, phylogenetic relationship, gene expression

## Abstract

Invertases and their inhibitor proteins are key regulators of carbon allocation in plants. Manipulation of invertase inhibitor (ITI) activity can potentially increase crop yield. The aim of this study was to determine the sequence diversity, phylogenetic relationships, and expression profiles of ITI genes in *sweetpotato*(*Ipomoea batatas*).. The coding sequences of two ITI paralogs (SPITI1 and SPITI2) were cloned from two sweetpotato varieties (Beauregard and Jewel) and sequenced. The DNA sequences were used to deduce amino acids sequences and predicted protein properties. Quantitative PCR (qPCR) was carried out to study the expression profiles of the genes at different developmental stages. The results show that introns are absent in both SPITI paralogs. SNPs, Indels, and variable simple sequence repeats (SSR) were present in the SPITI1 paralog, however, only SNPs were identified in the SPITI2 paralog. The predicted SPITI1 protein had 168, 172, or 174 amino acid residues, and molecular weights ranging from 17.88 to 18.38 kDa. In contrast, SPITI2 coded for a protein with 192 amino acid residues, with molecular weight ranging from 20.59 to 20.65 kDa. All conserved domains of ITI proteins were present in both protein isoforms.

Phylogenetic analysis indicated that SPITI genes were more closely related to *I.trifida* and *I.triloba* than *I.nil*, thus, suggesting their evolutionary relationship and conservation. A qPCR study indicated that both *SPITI* genes were expressed in all the sample tissues, though relative expression values differed across tissues at different developmental stages. This is the first study reporting diversity of *SPITI* genes and of an ~18 kDA isoform in sweetpotato. The findings may enable design of genetic engineering strategies for *SPITI* genes, including CRISPR/Cas gene editing in sweetpotato.

## INTRODUCTION

Sweetpotato is rated as the seventh most important food crop globally (Wu, et al., 2018). In China, the crop is rated as the fourth most important food crop (Yang, et al., 2017). In the US, sweetpotatoes have become more popular in recent years with consumption increasing nearly 42% between 2000 and 2016, and reaching 7.2 pounds per capita (North Carolina State University, 2018). Sweetpotato is also becoming increasingly important in Africa. There is a growing consumer demand which may be attributed to the promotion of the storage roots health benefits. In particular, consumption of orange-fleshed sweetpotato varieties are being encouraged in many African countries to promote health (Kofi Anan Foundation, 2021). Besides human consumption, sweetpotatoes are important for animal feed. In addition, the high energy content, and low requirements for cultivation makes the crop a potential raw material for bioenergy production (Lareo & Ferrari, 2019). As the crop becomes more important, different biotechnology approaches are needed for genetic improvement of the crop to meet the increasing demand, improve quality, and improve adaptability.

Regulation of invertase activity is one key route to improve crop yield. There are three types of invertases in plants, based on their subcellular location. These are vacuolar invertases, apoplastic (cell wall) invertases, and cytoplasmic invertases (Sturm, 1999). Cell wall invertase (CWI) irreversibly cleaves sucrose into glucose and fructose to enable apoplastic loading and unloading of sucrose into the phloem:companion cell complex (Xu, Hu, Yang, & Jin, 2017; Ruan, Jin, Yang, Li, & Boyer, 2010; Braun, Wang, & Ruan, 2014). CWIs determine the apoplastic sucrose:glucose ratio and regulate related signaling pathways. CWIs also maintain sucrose concentration gradients for sugar transport, storage, and partitioning between source and sink tissues (Chourey, Jain, Li, & Carlson, 2006).

CWI activity is highly regulated at both the transcriptional and post-transcriptional levels (Huang G.-J., et al., 2008). Post-translational activity of CWIs is controlled by a proteinaceous inhibitor called invertase inhibitor (ITI) (Krausgrill, Sander, Greiner, Weil, & Rausch, 1996).

ITIs are small molecular weight proteins which function to inhibit invertases by competitive inhibition. They bind to the active site of sucrose, thereby preventing invertases from binding to sucrose for cleavage (Rausch & Greiner, 2004; Zhang, Jiang, Yang, & Wang, 2015). Regulation of ITI activity indirectly enhances invertase activity, resulting in higher yields. Down-regulation of an ITI gene in tomato led to increase in seed weight, increase in fruit hexose level, and a prolonged leaf life span (Jin, Ni, & Ruan, 2009). Divora et al. (Sederoff lab; unpublished) observed that antisense repression of *ITI* in *Camelina sativa* resulted in shorter internodes (hence shorter plants) than the wild type control plants, although the silenced plants had higher biomass and seed yield.

*ITI* genes have not been well studied in sweetpotatoes. To-date, only one sequence (SPITI) from the species has been cloned and characterized. The first and the only study was by Huang et al, (2008) who characterized one *SPITI* homologue from the storage root. They reported that *ITI* from sweetpotato codes for a 192 amino acid protein with a predicted molecular weight of 20.624 kDa. Adequate knowledge of ITI genes in sweetpotato, including sequence diversity, phylogenetic relationships, and expression pattern is needed to enhance design of appropriate regulatory technologies, including CRISPR mediated gene editing of ITIs genes in the crop. Thus, the aim of this study was to characterize ITIs genes in sweetpotatoes, determine their phylogenetic relationships and expression profiles. We report that in addition to the 192 aa residue isoform, SPITI isoforms of 168, 172 and 174 aa residues with predicted Mw of 17.88 to 18.6 kDa are also present in sweetpotato. We also show that ITI in sweetpotatoes are more closely related to ITIs in *I.trifida* and *I.triloba* than they are to *I. nil*, thus, highlighting the phylogenetic relatedness among sweetpotatoes and sweetpotato wild relatives. The expression pattern of the two paralogs are also presented.

## MATERIALS AND METHODS

### Plant Materials

Two sweetpotato varieties, Beauregard and Jewel, were used in this study. Beauregard is one of the most popular sweetpotato varieties in the United States (Rolston, et al., 1987). It has long internodes and twinning tendency, with the vines reaching over 3 to 6 feet in length. It has fast growth and matures relatively early in 105-110 days. The variety has very high yields. The Jewel (NC Agricultural Research Service, 1971), now considered somewhat of an heirloom cultivar, is also one of the common sweetpotato varieties in breeding programs across the globe (Pope, Nielsen, & Miller, 1971). Unlike the Beauregard, Jewel has slow rate of growth, shorter internodes and no twinning tendency. Jewel needs about 120-135 days growing time for maximum yield (Scripps Networks, LLC., 2019). The two are orange-fleshed varieties that are used in many breeding programs. In vitro virus-tested materials were obtained from the Micropropagation and Plant Repository Unit (MPRU) at NC State University, and were kept in vitro by regular subculture.

### Cloning of SPITI genes

Genomic DNA was extracted from young leaves using a Qiagen DNeasy Plant Mini Kit (QIAGEN Inc., Germantown, USA). The manufacturer protocol was followed to extract DNA from the two sweetpotato varieties. The quality and quantity of the DNA were assessed using the NanoDrop (Themofisher, Waltham, Massachusetts, USA). The DNA was then stored at −20°C until further processing. To amplify the two *ITI* genes from sweetpotato, *ITI* sequences identified from NCBI database (from *Ipomoea nil* LOC109175922 and LOC109170182) were used as queries in the Sweetpotato Genomics Resource (http://sweetpotato.uga.edu/) to identify *ITI* homologues in *I. trifida and I. triloba*. Primer pairs (CWII_P18F + CWII_p18R; and CWII_P20F + CWII_P20R) (Table 1) were designed from conserved regions in the sequences, with the aid of online IDT Oligo Analyzer Tool (Integrated DNA Technologies, Inc., Coralville, USA) and NEB Tm Calculator (New England Biolabs Inc., Ipswich, Massachusetts, USA) to amplify the full-length coding region. PCR amplification was carried out using NEB Q5 High Fidelity 2x Master Mix (New England Biolabs Inc., Ipswich, Massachusetts, USA). The PCR reaction consisted of 2.5 ul each of 10 μM Forward and Reverse Primers; 25 μl of Q5 High-Fidelity 2X Master Mix; 900 ng of genomic DNA; and nuclease free water to a final volume of 50 ul. The cycling conditions were as follows: Initial denaturation for 30 seconds at 98°C; followed by 35 cycles of 98 °C for 10 seconds; annealing temperature at 64 °C; and 72°C for 20 seconds extension. The final extension was 72°C for 2 minutes, and a hold at 4°C. The PCR product was run on 1% agarose gel containing 1% ethidium bromide in TAE buffer. The gel was visualized under UV illuminator. The expected bands were cut, purified and cloned into PGEM-T Easy Vector (Promega, USA) using the manufacturer’s protocols. The ligated plasmid was used to transform *Escherichia coli* cells. The transformed cells were plated on LB plates containing 50 mg/l Carbenicillin, Isopropyl β-D1-thiogalactopyranoside (IPTG), and 5-Bromo-4-chloro-3-indolyl β-D-galactopyranoside (X-Gal) for blue white selection (Gold Biotechnology, 2016).

**Table 1:**
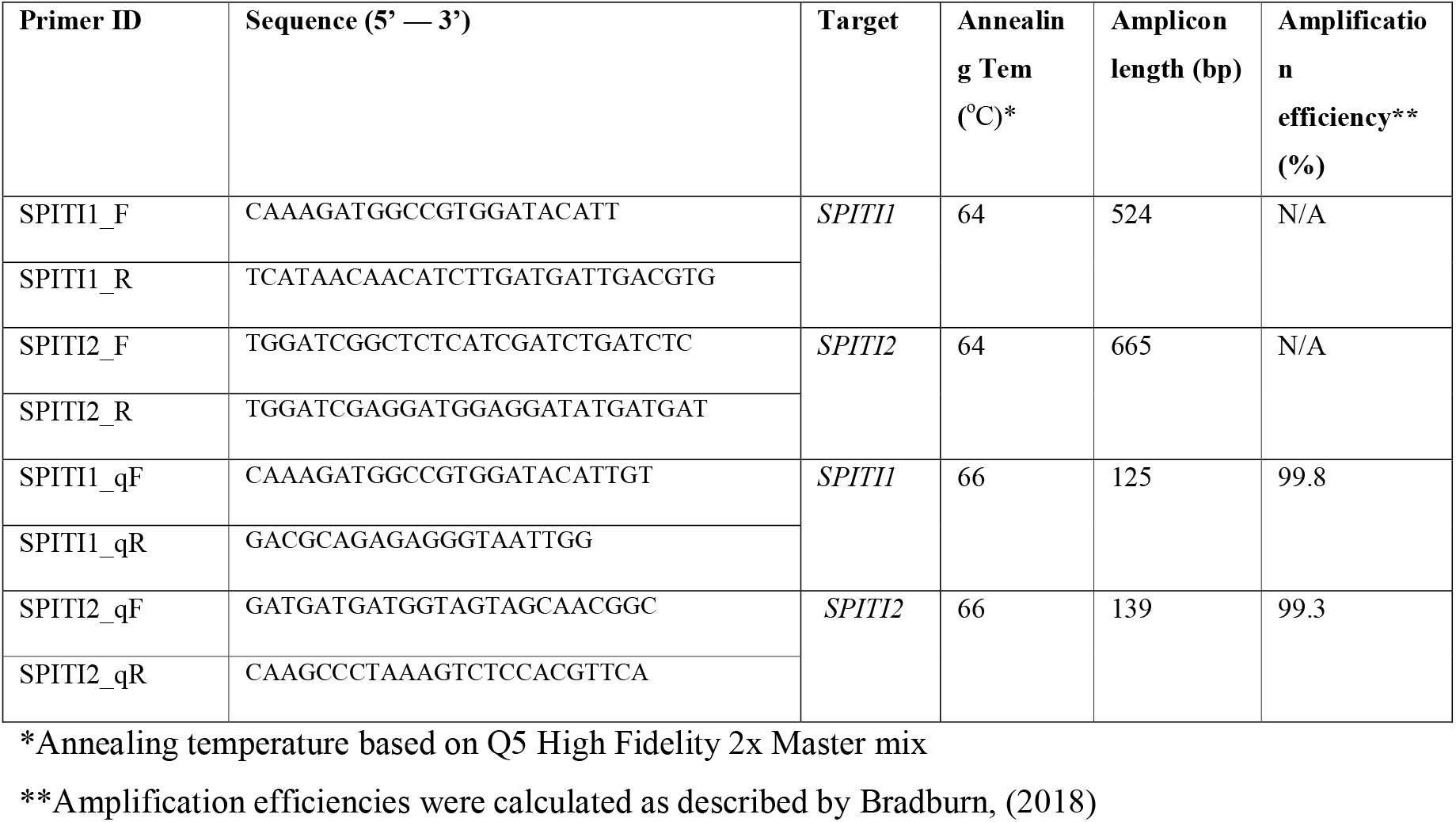
Primers used for cloning and qPCR studies of ITI genes in sweetpotatoes

### DNA sequencing

Twenty positive colonies were randomly picked for each gene per variety for plasmid purification. Purified plasmids were sent for Sanger Sequencing at GENEWIZ (Research Triangle Park, NC., USA).

### In silico characterization of ITI proteins in sweetpotato

Different online tools were employed to characterize the sweetpotato ITI genes and ITI proteins. Protein sequences were predicted from DNA sequences using the online ExPAsy translate tool (https://web.expasy.org/translate/) and NCBI blastx. Multiple sequence alignment was conducted using the online tools MUSCLE (Multiple Sequence Comparison by Log-Expectation) alignment (https://www.ebi.ac.uk/Tools/msa/muscle/) and NCBI nucleotide BLAST (Altschul, Gish, Miller, Myers, & Lipman, 1990). Phylogenetic trees were constructed using Molecular Evolutionary Genetics Analysis (MEGA) software package (https://www.megasoftware.net/).From the protein sequence, the isoelectric point (pI) and molecular weight (MW) of each protein were determined with the aid of Protein Molecular Weight (https://www.bioinformatics.org/sms/prot_mw.html) and ExPAsy Compute pI/Mw (https://web.expasy.org/compute_pi/). Protein motifs were determined with motif prediction tool MEME (http://meme-suite.org/index.html). Different programs, including Phobius (https://www.ebi.ac.uk/Tools/pfa/phobius/) (Madeira, et al., 2019), TargetP (Server 2.0; https://services.healthtech.dtu.dk/service.php?TargetP-2.0) (Emanuelsson, Nielsen, Brunak, & Heijne, 2000), and SignalP (Sever 5.0; http://www.cbs.dtu.dk/services/SignalP/ (Almagro Armenteros, et al., 2019) were used to predict presence or absence of signal peptide, cellular localization, and cleavage site, respectively. GC content was analyzed with the online tool Genomics %G~C Content Calculator (Science Buddies, https://www.sciencebuddies.org/science-fair-projects/references/genomics-g-c-content-calculator).

### Phylogenetic analysis

A phylogenetic tree was constructed to compare the structure of the predicted proteins with characterized ITI proteins from sweetpotato and other species. The phylogenetic tree was constructed with the neighbor joining method in MEGA X (https://www.megasoftware.net/).

### Characterization of gene expression by qPCR

*In vitro* plantlets of Beauregard and Jewel were acclimatized to greenhouse conditions for four weeks. The young plants were then transferred into soil medium in larger 15 cm pots. The plants were maintained at a temperature of 28-32 °C and 60-90% humidity. The plants were irrigated as needed. Sample plants were harvested for RNA extraction from week 3 to week 15. Sample tissues were collected from three plants for each variety. From each plant, three fully expanded mature leaves were collected from the upper third of the plant and pooled per sample. The leaves, roots and storage roots were collected using the same pooling procedure. The plant tissues were collected in liquid nitrogen, ground and stored at −80 °C until RNA extraction.

RNA extraction was carried out using Qiagen RNeasy kit (Qiagen, Germantown, MD, USA). RNA concentration was determined using NanoDrop and the integrity was assessed by gel electrophoresis. Aliquots of 2.0 ug of the total RNA were collected for cDNA synthesis by using Clontech Oligo dT cDNA synthesis kit (Takara Bio USA, Inc., USA following the manufacturers protocol.

To study the expression of the two genes, the Sanger sequences were aligned for each gene paralog to identify conserved regions in mRNA sequences. qPCR primers were then designed from the conserved regions with the aid of IDT OligoAnalyzer Tool (https://eu.idtdna.com/pages/tools/oligoanalyzer) and NEB Tm Calculator (https://tmcalculator.neb.com/#!/main). qPCR conditions were optimized for each pair of primers. Different sets of primers were evaluated, however, the set of primers with the best amplification efficiency for each gene was used for the qPCR study and are reported in Table 1. The cycling conditions were based on the Q5 high fidelity 2x Master Mix reaction protocol. The qPCR reaction conditions were as follows: Initial denaturation at 98 °C for 30 seconds; followed by 40 cycles of 98°C for 10 seconds, 64°C for 20 seconds.

### Experimental design and data analysis

The expression study followed a complete randomized design. Data were taken from three biological replicates, and at least two technical replicates. Expression levels were normalized using TUB, H2B and ACT as internal control genes. Data were analyzed using ANOVA, Mean differences were considered significant when p-values were less than 0.05. Means were separated using Tukey Test.

## RESULTS

### DNA sequence diversity among ITI genes in sweetpotato

PCR was carried out, using genomic DNA as a template, to amplify ITI genes in Beauregard and Jewel varieties. The band size (about 540 bp and 670 bp for SPITI1 and SPITI2 paralogs, respectively) of the amplified product was the same as the expected band size from complementary DNA (cDNA). The above observation suggested the absence of intron(s) in the transcribed region of the gene. In order to confirm this, PCR was conducted on cDNA to amplify the target sequence, and resolved on agarose gel, alongside PCR product from genomic DNA. Figure 1 provides the gel image for SPITI1 homologue. From the gel image, it can be observed that the band sizes in all the sample lanes are the same (about 540 bp), irrespective of the PCR template, indicating absence of intron(s) in the transcribed region of SPITI1 gene in the sweetpotato genotypes. Similar observation was made on the gel of SPITI2 paralog. No intron was observed.

**Figure 1:**
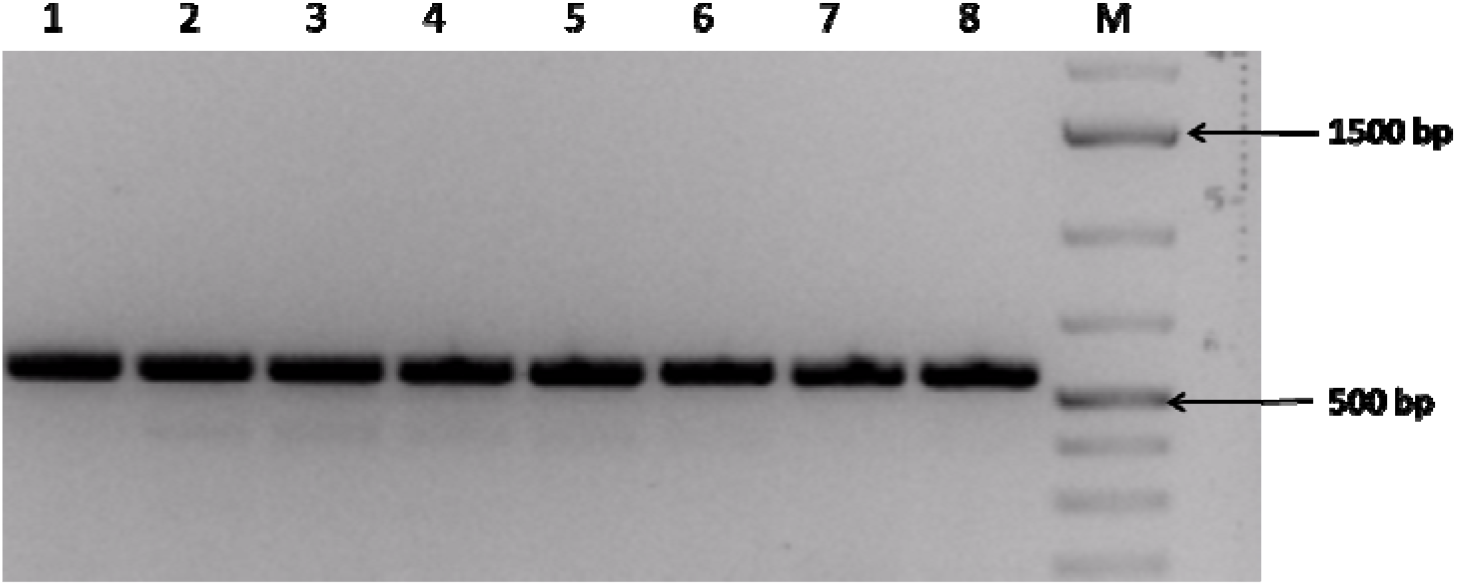
PCR amplified product of CWIIA gene homologue in sweetpotatoes resolved on agarose gel. PCR and gel electrophoresis were performed with two technical replicates. Lanes 1 and 2: products of genomic DNA of Beauregard; 3 and 4: product of cDNA of Beauregard; 5 and 6: genomic DNA of Jewel; 7 and 8: cDNA of Jewel. M is DNA ladder (Thermo Scientific™ GeneRuler 1 kb Plus DNA Ladder)

To determine the nucleotide sequence, the PCR product from genomic DNA of the two sweetpotato varieties were cloned, and sequenced. The sequences were submitted to the National Center for Biotechnology Information (U.S. National Library of Medicine, Bethesda MD, 20894, USA) and were assigned accession numbers as presented in Table 2. In all, 15 alleles were identified from the two sweetpotato cultivars; six (6) alleles for the SPITI1 paralog, and nine (9) alleles for the SPITI2 paralogue. The coding sequence of all SPITI2 alleles were 579 bp long. In contrast, the coding sequence of SPITI1 varied between 507 bp, 519 bp, and 525 bp.

**Table 2:**
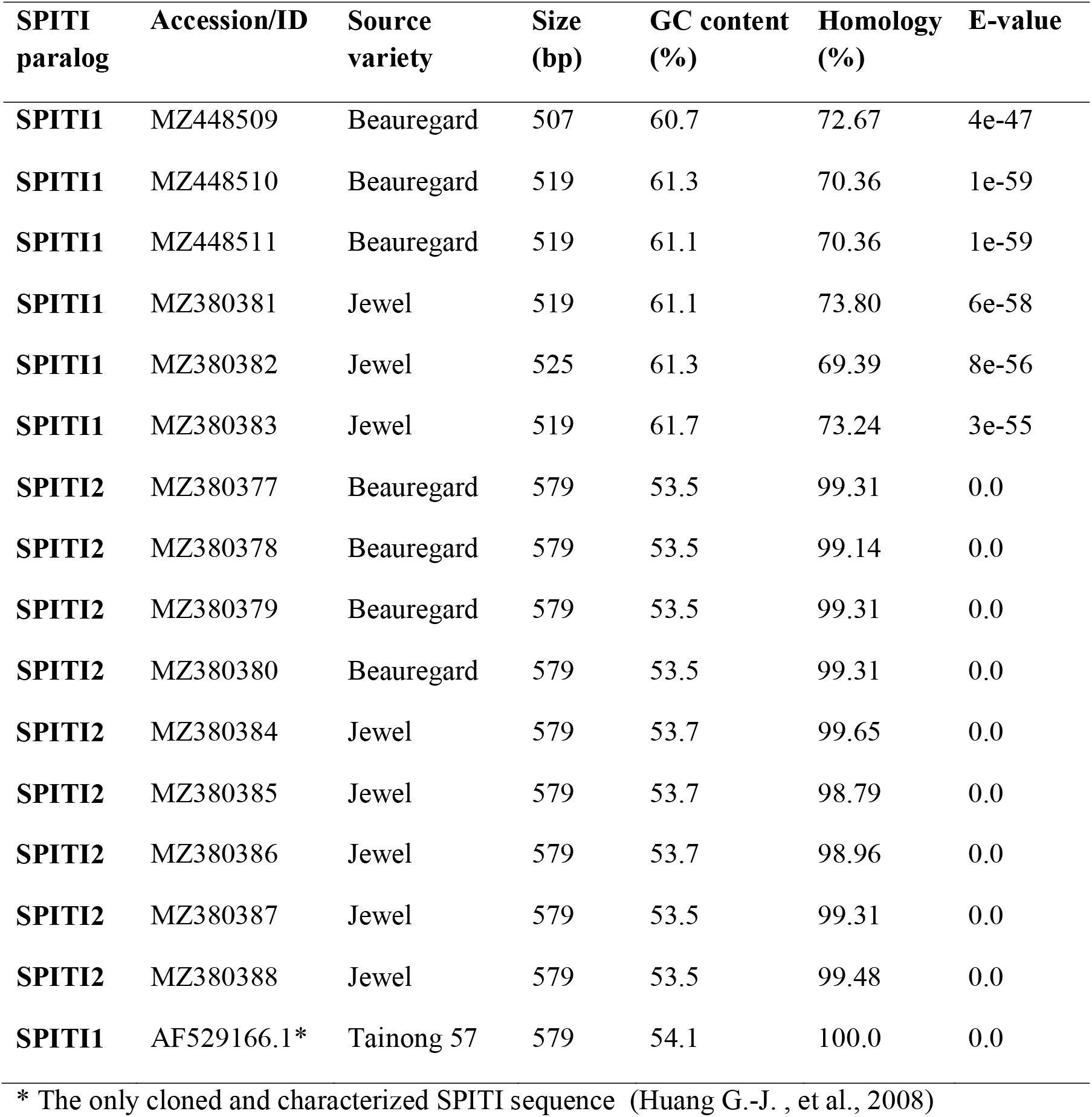
Characteristics of invertase inhibitor genes in sweet potatoes

For a more effective analysis and comparisons, sequences from *Arabidopsis thaliana*, tomato (*Solanum lycopersicum*), potato (*Solanum tuberosum*), and *Camelina sativa* were retrieved from NCBI nucleotide database and used as outgroups. ITIs from potato, tomato, *Arabidopsis*, and *Camelina* were selected because they are among the best studied and characterized invertase inhibitors. All the DNA sequences were trimmed to the open reading frame before being used for any analysis and comparison.

### GC content and sequence homology

GC content was significantly higher (p < 0.05), averaging 61.2% in SPITI1, than SPITI2 which had 53.6% average GC content. Among the *Ipomoea* species, GC content was lowest in *I.nil*, while the highest was observed in cultivated sweetpotato *(I.batatas)*. The average GC content for both paralogs averaged 56.8%. Between sweetpotato varieties, no significant differences were observed. However, compared with other genera, GC content was significantly higher (p < 0.05) in the *Ipomoea* species than the other species whose average was 39.8%.

At the NCBI database (Altschul, Gish, Miller, Myers, & Lipman, 1990) the highest homology found was the sequence of mRNA AF529166.1, a SPITI sequence isolated from Tainong 57, a yellow fleshed sweetpotato cultivar in Taiwan (Huang G.-J., et al., 2008). There was 98.8% to 99.7% homology between SPITI2 genes, contrary to SPITI1 genes with 55.73-72.36% homology. All sequences with significant homology were members of the plant pectin methylesterase inhibitors (PMEIs) superfamily domain, and pectin methylesterase inhibitor related protein (PMEI-RP) family (Hothorn, Wolf, Aloy, Greiner, & Scheffzek, 2004).

Homology values for all the accessions are presented in Table 2. The coding sequences of SPITI1 gene homologue in sweetpotatoes consists of 507-525 bp, with a translated protein of 168-174 aa residues (Supplementary Table S1) while SPITI2 consists of 579 bp with a translated protein of 192 aa residues.

### Single nucleotide polymorphism, nucleotide insertions and deletions

Single nucleotide polymorphisms (SNPs) were observed in both gene paralogs. In addition, DNA polymorphisms involving Indels (insertion and deletions), and substitutions were observed. Nucleotide deletions, nucleotide insertions and nucleotide substitutions were all present in the SPITI1 homologue. However, only nucleotide substitutions were observed in the SPITI2 homologue (Table 3). The number of polymorphic sites varied between the two paralogs. A total of 12 SNP loci were identified in the A homologue, representing 2.4 SNP loci per 100 bp. In the B homolog, six (6) SNP loci were observed, representing 1.03 SNP loci per 100 bp. All the SNPs led to amino acid substitutions. None resulted in premature stop codon. No frame shift Indels was observed.

**Table 3:** Diversity of SPITI genes in Beauregard and Jewel

| **Gene homologue** | **All paralogs** | **Homologue A** | **Homologue B** |
| --- | --- | --- | --- |
| Number of alleles | 15 | 6 | 9 |
| Number of nucleotides | 507-579 | 507-525 | 579 |
| Nucleotide substitutions | Present | Present | Present |
| Nucleotide Indels | Present | Present | Absent |
| Number of SNP loci | 149 | 12 | 6 |
| Number of SNPs loci per 100 bp | 25.73 | 2.40 | 1.03 |
| Number of Haplotypes (h) | 12 | 6 | 6 |
| Nucleotide diversity ($\pi$) | 0.12604 | 0.01087 | 0.00384 |
| Haplotype diversity (Hd) | 0.943 | 0.01087 | 0.833 |

On the whole, more sequence diversity was observed in the SPITI1 homologue, compared to the SPITI2 homologue. Nucleotide diversity (π) was higher (0.01087) in the A homologue than in the B homologue 0.00384. Similarly, haplotype diversity was higher in homologue A (0.943) than homologue B (0.833).

Sections of the nucleotide alignment for homolog A are presented in Figure 2. The full DNA sequence alignment is shown in Supplementary File S1. Alignment for SIPITI genes is presented in Supplementary File S2. It can be observed from the alignment that a 12 nucleotide deletion is present in the allele MZ448509. The sequence GCGGCGGTGGAG is absent in that allele. However, that sequence is conserved in all other alleles, including sequences from *I. trifida* and *I. triloba*. The deletion, however, did not affect the open reading frame.

**Figure 2:**
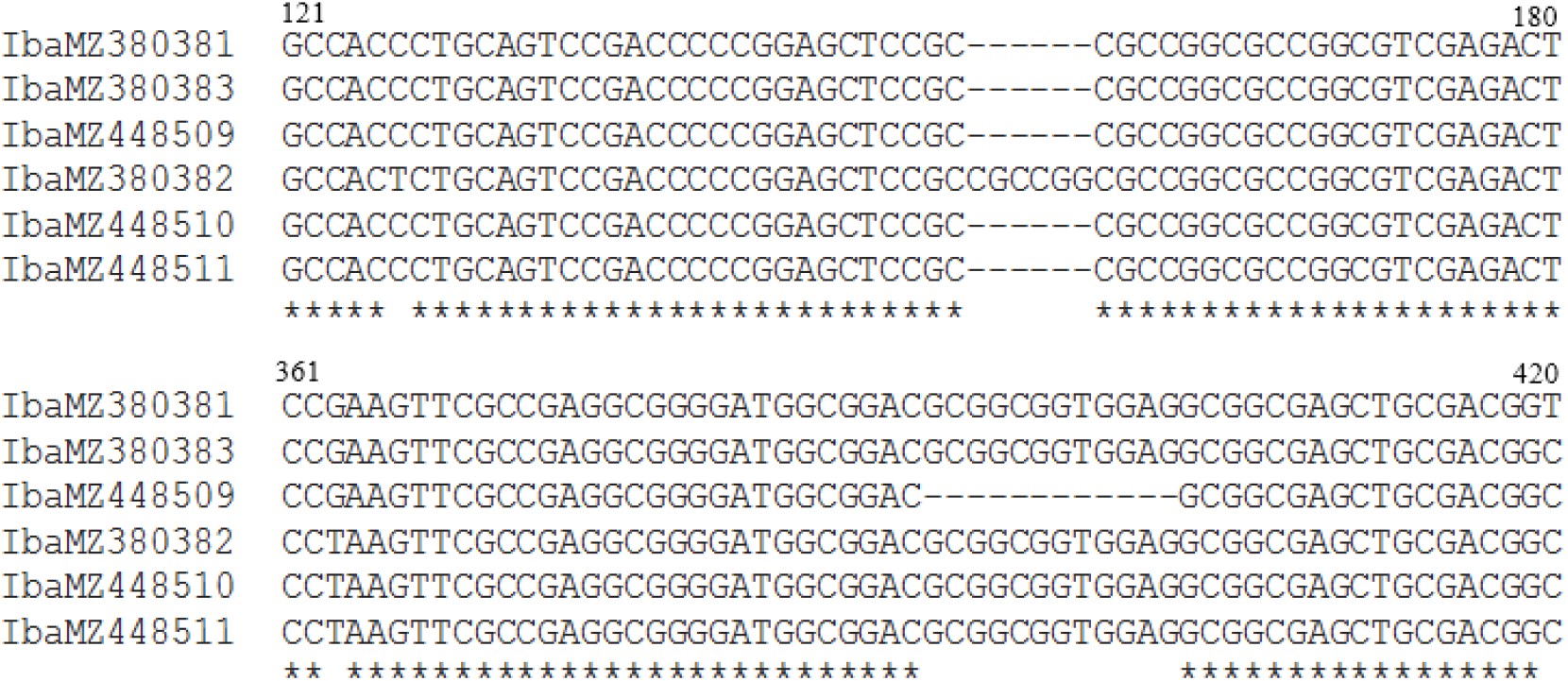
A section of nucleotide alignment of SPITI1 homologue in two sweetpotato varieties.

The sequence (CGCCGG)n is a hexanucleotide simple sequence repeat (Microsatellite) identified in the SPITI1 homologue. The sequence is however, absent in the SPITI2 homologue. In most of the SPITI1 alleles, the sequence is repeated two times (CGCCGG)_2_. However, in the MZ380382 allele, the sequence is repeated three times (CGCCGG)_3_, thus, representing a trinucleotide substitution in that sequence. In *Arabidopsis* and *I.nil*, the sequence is present only once. The sequence is absent in tomato and potato homologues.

Within the SPITI2 homolog, another microsatellite (SSR) is observed. A trinucleotide tandem repeat (CTA)_n_ is present in the sweetpotato homologue. The sequence is present three times (CTA)_3_ in majority of the alleles. However, in the itb01g35980.t1 allele, it is present four times (CTA)_4_, thus representing three nucleotide insertion (one amino acid insertion).

Thus, the itb01g35980.t1 allele translates into a 193 aa residues protein, compared to the rest of the alleles which all translated 192 aa proteins (Supplementary Table S1).

### Diversity of ITI proteins in sweetpotatoes

The protein sequence alignmen When the sequences were compared with AAM94391.1, the characterized SPITI protein in the *Ipomoea* genus, sequence homology was between 54.21% and 55.79%, with an average of 55.34±0.618%. Within the wild relatives, homology was between 55.21% and 55.73% with an average of 55.480±0.261%. Homology was significantly higher (p<0.05) in the SPITI2 isoform as compared to the SPITI1. Though no significant difference was observed within the *Ipomoea* genus, homology was higher in cultivated sweetpotato (average 98.560±0.14%) compared to the wild relatives (average 94.30±3.6%). However, when compared with non *Ipomoea* species, homology was significantly very low (average 42.85±3.67%). All expected values (E-values) were significant.

Isoelectric point values significantly differed between the two protein isoforms (Supplementary Table S1). pI values ranged between 4.69 and 5.13 with an average of 4.97±0.16 in the SPITI1, while pI range was between 4.45 and 4.47, with an average value of 4.46±0.011 in the SPITI2. No significant difference was observed between the average pI values of cultivated sweetpotatoes and the wild relatives. Also, no significant differences were observed between alleles from Beauregard and Jewel. The highest pI in *Ipomoea* was 5.13. When compared with other genera, pI values of the *Ipomoea* Genus was significantly lower.

### Secondary structure and Motifs present in SPITI proteins

Conserved motifs were identified with the online motif prediction tool MEME (http://meme-suite.org/index.html). For comparison and accurate analysis of SPITI proteins, the amino acid sequences from other species *(I.nil, I.trifida*, and *I.triloba)*, potato *(Solanum tuberosum*, tomato (*Solanum lycopersicum*), *Arabidopsis thaliana* and *Camelina sativa* were added in the analysis. Output from the MEME program is presented in Figure 3. Out of the 14 motifs, seven were present in the SPITI1 isoforms in *Ipomoea spp*, with the exception of MZ448509 which lacked motif 7. Motifs absent in SPITI1 were 5, 9, 10, 13, and 14. In contrast to the SPITI1, eight conserved motifs were present in the SPITI2 isoform. Motifs absent in the B isoform included motifs 8, 10, 13, and 14. Motifs 10, 13, and 14 are absent in *Ipomoea spp*. Motifs 5, 8 and 9 were unique to the Genus. The functions of the motifs have not been determined.

**Figure 3:**
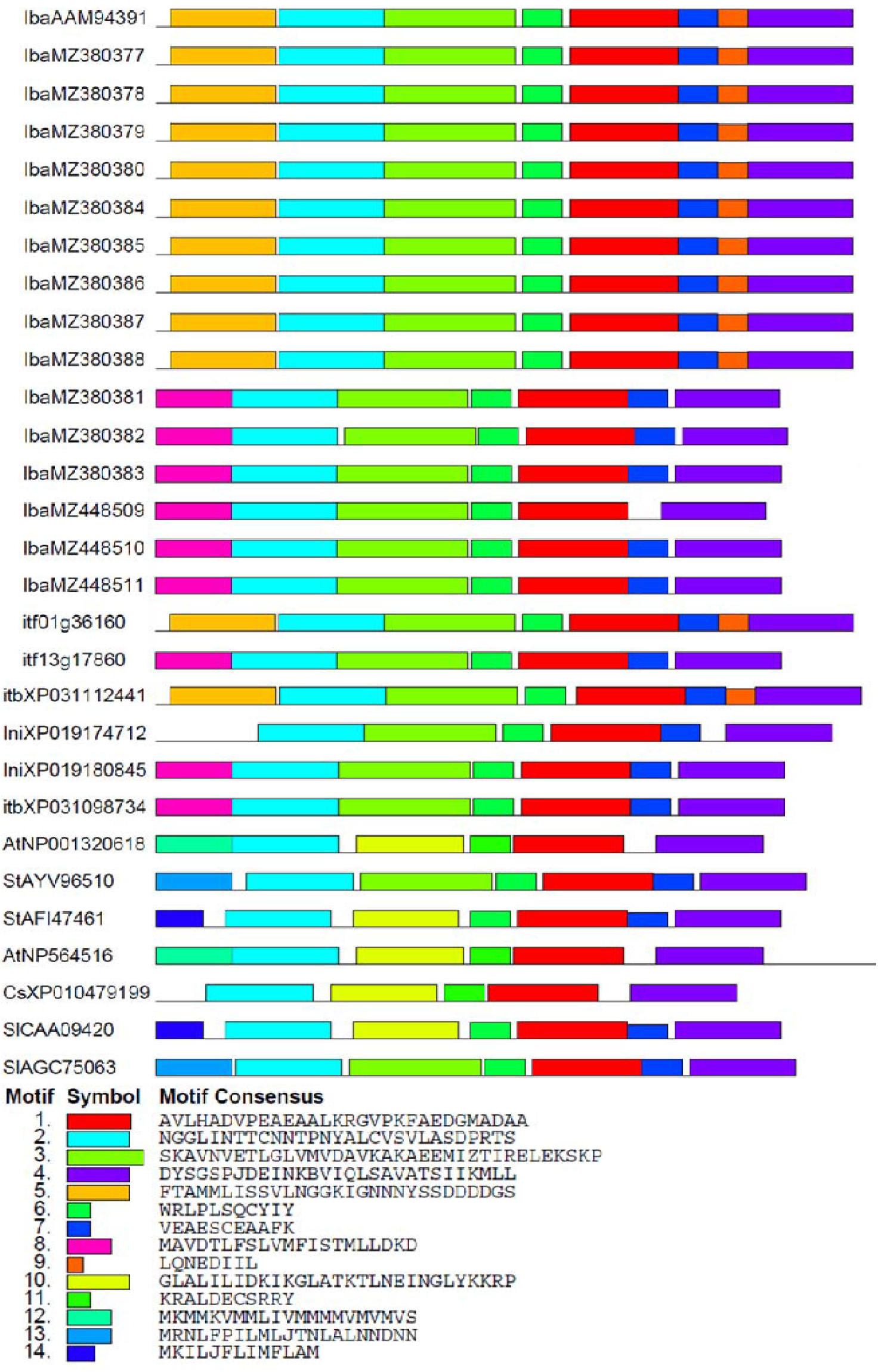
Conserved motifs deduced from SPITI proteins in sweetpotatoes. Accession numbers are preceded by 2 or 3-letters to identify the species. Iba, *Ipomoea batatas;* Itf, *Ipomoea trifida;* Itb, *Ipomoea triloba;* Sl, *Solanum lycopersicum;* Csa, *Camelina sativa;* Ath, *Arabidopsis thaliana*.

The sequence and the set of homologs detected by PSI-Blast were processed by Phyre2 (http://www.sbg.bio.ic.ac.uk/phyre2/html/help.cgi?id=help/interpret_normal) to predict secondary structure, family, the presence of transmembrane helices, and to determine topology of the helices in the membranes. The secondary structures were predominantly α-helix (5), coil, with no β-strands. For the SPITI1, a total of seven Alpha helices, covering 84% and eight coils (disordered) covering 16% of the protein sequence were predicted. In the SPITI2 too, the secondary structure was predominantly alpha helix. However, the percentage of alpha helix was lower (74%) with 9% of the helix being transmembrane. A total of eight alpha helices were identified in the SPITI2 isoform.

### Sub-cellular localization

A signal peptide was predicted in Isoform SPITI2 (Figure 4) from aa residues 1-19. A significant portion of the protein lie within the extra cellular space (apoplast), with aa residue 173-189 embedded within the membrane, while the C-Terminal end lie within the cytoplasm. No signal peptide was predicted for any of the SPITI1 isoforms.

**Figure 4:**
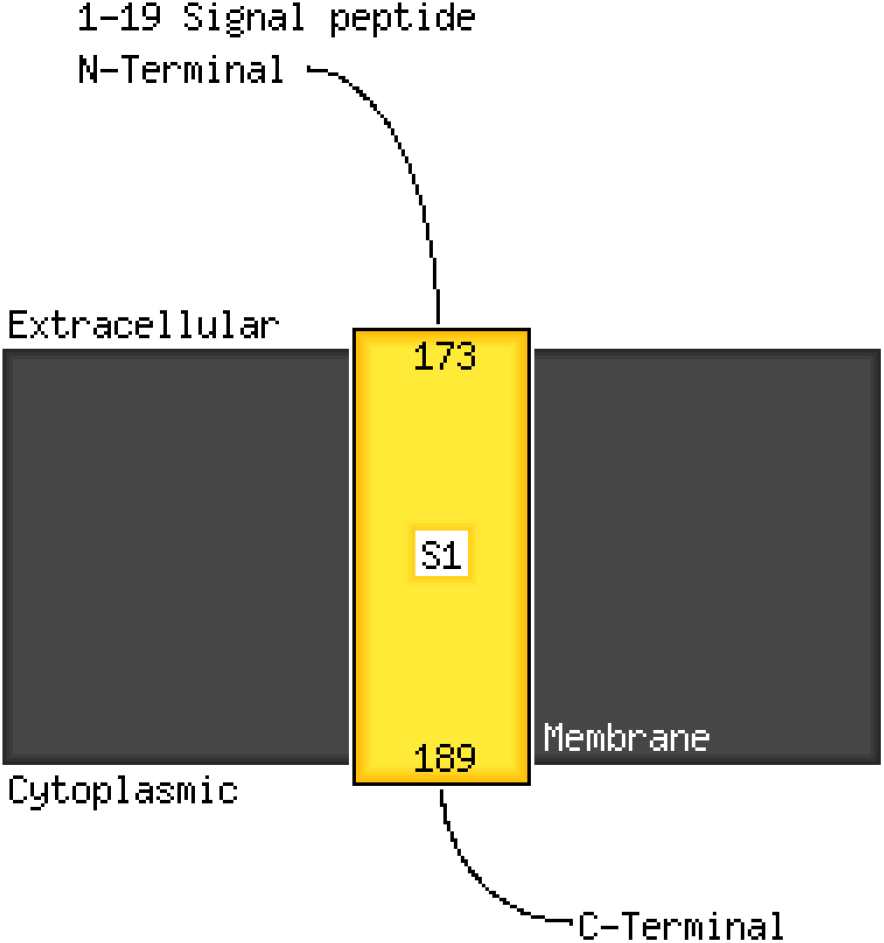
Predicted signal peptide and topology of SPITI2 protein in the membrane.

The predictions and associated probabilities as well as cleavage position predicted by SignalP and other programs are presented in Supplementary Table S2. No signal peptide was predicted in any of the predicted proteins of SPITI1. In contrast, a signal peptide was predicted in all the predicted proteins of SPITI2. All the predictions values were high, and associated with high probabilities. The position of cleavage was the same for all the predicted proteins, (except predicted protein of the allele MZ380386) and occurred between amino acids 17 and 18. In the MZ380386 however, cleavage position occurred at aa 21 and 22.

SPITI1 proteins were generally predicted to have widespread localization in the cell, with little or no localization in the apoplast (cell wall). Localization in the nucleus, vacuoles, and ER also were predicted. In contrast, the SPITI2 isoform was predicted to be highly localized within the apoplast, followed by the chloroplast. There was no predicted localization in the vacuoles, the nucleus, and the ER.

### Phylogenetic Relationships among predicted *ITIs* proteins from sweetpotato, and other species

There was sequence conservation at both the DNA and amino acid level. The protein sequence alignment and phylogenetic tree are presented in Supplementary File S3. All common domains typical of cell wall invertase inhibitor proteins were present in the predicted proteins. These include four conserved cysteine residues, and a PKF motif.

The phylogenetic tree generated using amino acid sequence of SPITI proteins from the current study is presented in Figure 5. Sequences from sweetpotato wild relatives, *I.nil, I.trifida* and *I.triloba* as well as sequences from *Arabidopsis* and *Solanum* were included as outgroups. From the tree, two major groups (A and B) can be identified. Clade A, comprises proteins from *Ipomoea* spp (cultivated sweetpotato and sweetpotato wild relatives) while Clade B, comprises proteins from species other than *Ipomoea*. These were sequences from tomato (*Solanum lycopersicum)*, potato (*Solanum tuberosum*), *Arabidopsis*, and *Camelina*. Within the B clade, the proteins were grouped based on species, indicating protein conservation in species.

**Figure 5:**
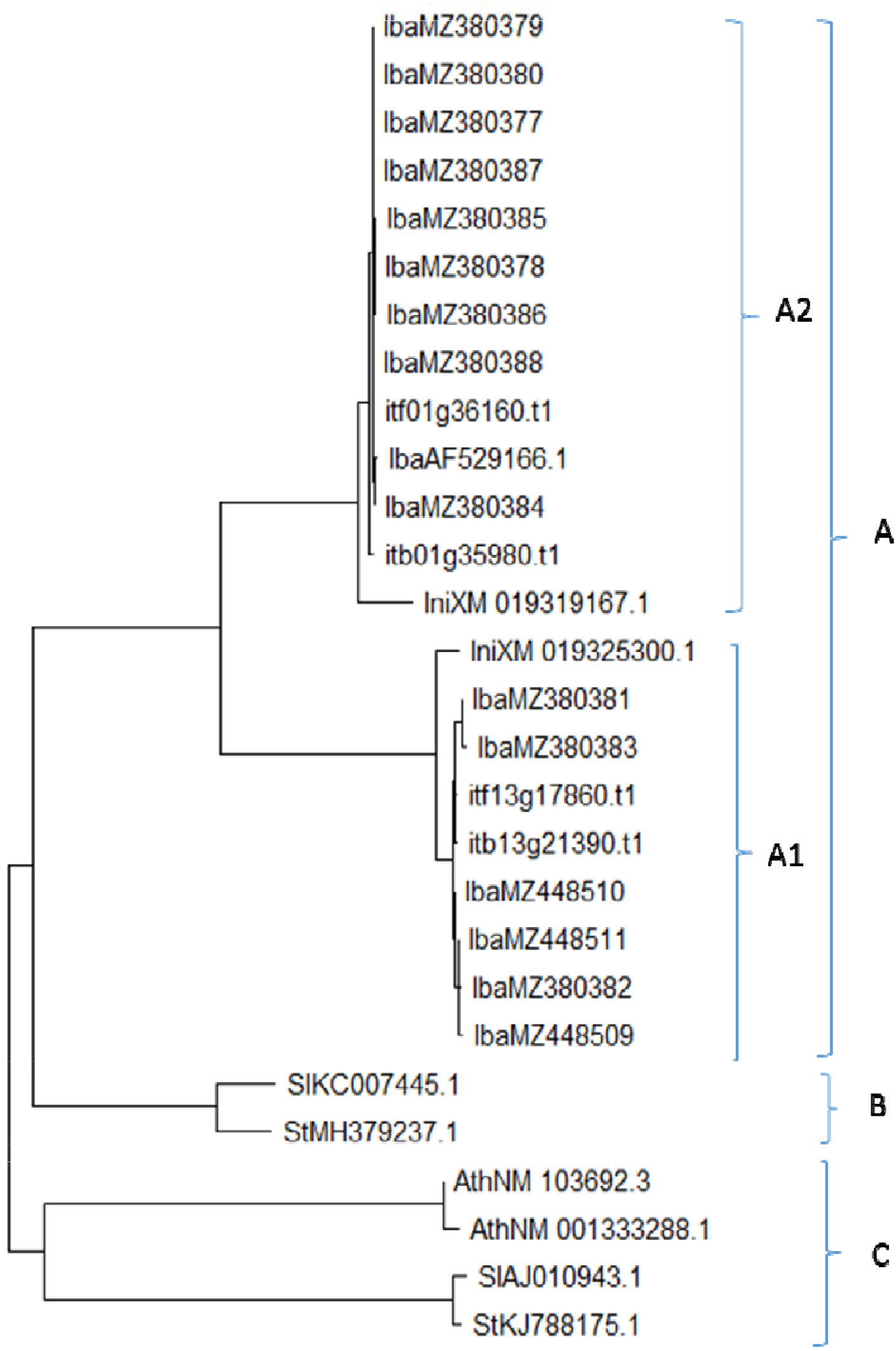
Phylogenetic relationships among ITIs genes in sweetpotato and other species.

The A clade comprises all sequences from *Ipomoea* species. Within the Clade A, two sub-group can be distinguished, A1 and A2, corresponding to SPITI1 and SPITI2 proteins respectively. AAM94391.1 was more closely related to MZ380384, a sequence found in the Jewel cultivar. The two paralogs share about 99.65 sequence homology. Within the *Ipomoea* clades A1 and A2, ITI protein from cultivated sweetpotato were grouped more closely to ITI protein from *I.trifida* followed by *I.triloba* while sequences from *I.nil* were further apart.

### Expression analysis of *SPITI* gene paralogs in sweetpotato

Relative expression of the two *SPITI* paralogs was observed to differ across tissues and developmental stages. Both paralogs were expressed in all the sample tissues studied. However, the expression pattern of the two paralogs were different, especially in the storage roots and the leaves.

#### Expression of SPITI1 paralog

Figure 6 shows the expression profile of SPITI1 paralog in Beauregard and Jewel. In the Beauregard, highest expression was observed in the leaves during the early plant growth stages (up to week 4). There was high expression in the storage roots as the storage roots began forming. However, relative expression reduced significantly in the storage roots, as the plant matured. High expression was observed in the roots, especially at week 8. Though relative expression levels fluctuated from one tissue to the other over the course of development, in general, highest expression was observed in the leaves.

**Figure 6:**
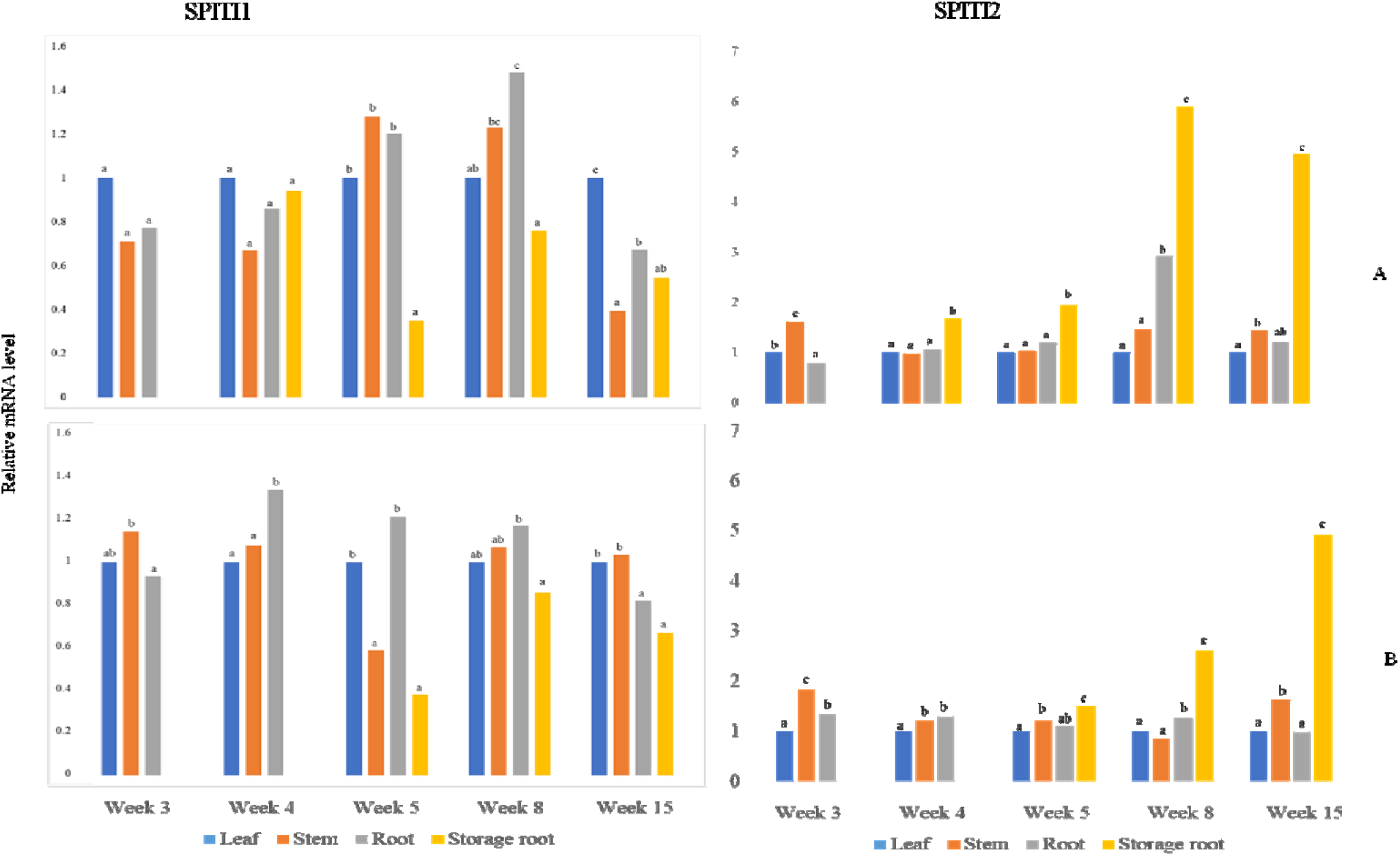
Expression profiles of SPITI genes in sweetpotatoes. Expression in Beauregard (A); and expression in Jewel (B). Means from the same week with different letters are significantly different at 0.05 level of significance.

In the Jewel variety, lowest expression was observed in the storage roots, while highest expression was in the roots. Relative expression fluctuated across developmental stages. In the Jewel, no storage roots formation was observed at Week 4. Storage roots were observed Week 5.

### Expression of SPITI2 paralog

The SPITI2 paralog was observed to be highly expressed in all the tissue samples studied. In the Beauregard, relative expression was highest in the storage roots, and lowest in the leaves. There was a general increase in rate of expression from the fourth week, as the storage roots accumulated starch. The relative expression level of SPITI2 declined at Week 15, though, the mean difference was not significant (p>0.05). At Week 8 the rate of expression in the storage roots was about six-fold, compared to expression in the leaves. However, relative expression was about fivefold at Week 15.

The expression pattern in the Jewel was similar to that of the Beauregard. In the Jewel, formation of storage roots was observed at Week 5. Relative expression in the storage roots increased consistently up to Week 15. While relative expression in the storage roots was about 1.5-fold in the storage roots as in the leaves, the fold increase was about five-fold at the maturity stage, Week 15.

## DISCUSSION

Cultivated sweetpotato is a hexaploid with 90 chromosomes, a high genome size, high diversity and high level of heterozygosity (Paliwal, et al., 2020; Karuri, et al., 2010; Yang, et al., 2017). Itt is thus expected that high diversity would be present in SPITI genes. Sequence information of ITI genes in sweetpotatoes may enable design of appropriate interventions to regulate the activity of the genes. In particular, design of guide RNA for the CRISPR/Cas system requires adequate knowledge of the sequence of the gene of interest (Jinek, et al., 2012). In this study, two paralogs of SPITI genes from two sweetpotato varieties were sequenced to enable design of appropriate intervention, including the CRISPR/Cas system for the genes. The study has provided sequences of 15 alleles for two SPITI paralogs in Beauregard and Jewel. The deduced amino acid sequences, secondary structures, and other properties of the proteins have been determined. The positions and DNA sequences for the four conserved cysteine residues which form two disulfide bridges to maintain the structure of the protein (Castrillon-Arbelaez & Delano-Frier, 2011; Su, et al., 2020) have been determined. The sequence of the PKF motif which targets the active site of the invertase, thereby blocking the invertase from functioning (Hothorn, Van den Ende, Lammens, Rybin, & Scheffzek, 2010) has also been determined. The sequence information may enable design of appropriate regulatory interventions, including genome editing and silencing of SPITI genes in the two sweetpotato cultivars.

Most reports have considered and treated ITIs as a single gene/single protein. In the current study, we observed that despite significant sequence homology, the two genes differ with regards to their rates of expression, expression breadth, and subcellular localization. They also differ in GC content, motifs, amino acid composition, and pI. The observed differences might suggest neofunctionalization or subfunctionalization of the paralogs following duplication of the gene in the species (Rastogi & Liberles, 2005).

The absence of introns in in both paralogs is consistent with absence of introns in the homologs from *I.trifida* and *I.triloba*. Absence of introns has also been observed in some invertase inhibitor homologs in potato (Brummell, et al., 2011), though introns are present in some homologs in potato (Datir, et al., 2012). In potato, the two paralogues are located on the same chromosome, and are separated apart by about 5.5 kb, with no gene in between (Datir, et al., 2012). In *I.trifida* and *I.triloba*. however, the two genes are located on different chromosomes.

The hexaploid genome of sweetpotato is believed to consist of three sub-genomes (B1B1B2B2B2B2) (Wu, et al., 2018). The first sub-genome (B1B1) is believed to have been derived from *I. triloba*. The second and third sub-genomes (B_2_B_2_B_2_B_2_) are believed to have been derived from a duplication of *I. trifida* genome (B_2_B_2_) (Wu, et al., 2018; Yang, et al., 2017; Isobe, Shirasawa, & Hirakawa, 2019). Sweetpotato genome is therefore considered both auto-and allo-hexaploid. It is thus expected that between two and six alleles for each gene be present in every sweetpotato variety. In this study, between three and five alleles were identified for each gene paralog, consistent with the genome structure of sweetpotato.

The 15 unique sequences from only two sweetpotato varieties imply high genetic diversity in sweetpotatocompared to other species. When Datir, et al., (2012) studied diversity of ITI genes in three tetraploid varieties of potato, a total of five alleles were identified (Datir, et al., 2012). In a similar study to compare diversity of ITI gene between processing and non-processing potato varieties, four alleles were identified from five potato varieties (Datir, Mirikar, & RaviKumar, 2019). Diversity within sweetpotatoes therefore, seems higher than what was observed in potato. The observed diversity is consistent with reported diversity in the sweetpotato genome. According to Yang et al, (2017), there are two or three polymorphic sites in every 100-150 bp in the sweetpotato genome. In the current study, an average of 2.4 polymorphic sites were present per 100 bp in the SPITI1, which is consistent with the above report. However, 1.03 polymorphic sites per 100 bp were observed in the SPITI2 homolog, implying a lower diversity in SPITI2.

Despite conservation in ITI genes, an appreciable level of diversity has been observed within certain regions of the gene. Some of the motifs present in sweetpotato and other *Ipomoea* species were unique to the genus. The phylogenetic tree (Figure 5) classified sequences in *Ipomoea* species into two distinct groups (A1 and A2), which corresponds to their homology, expression, and localization (Datir, et al., 2012). At the protein level, the two groups share about 55% homology. This, thus indicates that the two are non allelic, but rather encoded by different genes. Similar observations have been made in sugarcane (*Saccharum* spp) (Shivalingamurthy, et al., 2018), and potato (Datir & Ghosh, 2020).

The ancestry of cultivated sweetpotato has not been fully resolved. There are about 600 species in the *Ipomoea* genus, 13 of them are closely related to cultivated sweetpotato (Austin & Huaman, 1996). Studies involving cytology and molecular studies predicted *I. trifida* as the closest wild relative to sweetpotato. Recent studies utilizing next generation sequencing data have predicted *I. trifida* and *I. triloba* as the most closely related to sweetpotato, and therefore, the most likely progenitor of cultivated sweetpotato (Li, et al., 2019). Wu, et al., (2018) identified *I. trifida* as the closest relatives of sweetpotato. In the current study, the phylogenetic analysis indicated that sweetpotatoes are more closely related to *I. trifida*, followed by *I. triloba*, with *I.nil* being the least related. The current finding thus, agrees with previous studies which support the *I.trifida/I.triloba* ancestry of sweetpotato.

In both sweetpoato varieties, relative expression levels varied across different tissues at different developmental stages. Both paralogs had high expression in all the tissues studies. This observation is consistent with other studies reporting expression of ITI genes. In *Arabidopsis*, AtC/VIF1 and AtC/VIF2 displayed different target specificities and expression profiles (Link, Rausch, & Greiner, 2004). In potato, Brumell et al (2011) observed that ITI mRNA accumulated in all tissues, but at varying levels. They also reported differences in expression levels between the two homologs. In addition, they observed that one homolog accumulated at increasing abundance in the tubers as the tubers matured. In sweetpotato variety Tainong 57, Huang et al (2008) reported high levels of expression in the storage roots. They further observed lowest expression in fully expanded leaves. In the current study, mRNA levels of SPITI2 in both cultivars accumulated in the storage roots at increasing amounts as the storage roots matured, similar to the observation in potatoes.

There was relatively high expression of SPITI1 in the leaves of Beauregard, and this might require further investigation. The high accumulation of SPITI2 in the storage roots may imply an important role of the gene in the development of the storage roots. Thus, regulating the amount of expression in the storage roots might have significant impact on the development of storage roots and tubers. Technologies involving CRISPR-mediated gene editing, and RNA interference may help to evaluate the activity of the genes in the storage roots.

## ACKNOWLEDGEMENT

This study was funded by the Borlaug Higher Education for Agricultural Research and Development (BHEARD)/USAID at Michigan State University

## REFERENCES

Isobe, S., Shirasawa, K., & Hirakawa, H. (2019). Current status in whole genome sequencing and analysis of Ipomoea spp. Plant Cell Report, 38, 1365–1371. doi:https://doi.org/10.1007/s00299-019-02464-4

Almagro Armenteros, J. J., Tsirigos, K. D., Sønderby, C. K., Petersen, T. N., Winther, W. O., Brunak, S.,… Nielsen, H. (2019). SignalP 5.0 improves signal peptide predictions using deep neural networks. Nature Biotechnology, 420–423. doi:10.1038/s41587-019-0036-z

Altschul, S. F., Gish, W., Miller, W., Myers, E. W., & Lipman, D. G. (1990). Basic local alignment search tool. Journal of Molecular Biology, 215, 403–410.

Austin, D. F., & Huaman, Z. (1996). A synopsis of Ipomoea (Convolvulaceae) in the Americas. Taxon, 45, 3–38.

Baldwin, S. J., Dodds, K. G., Auvray, R. A., Macknight, R. C., & Jacobs, J. M. (2011). Association mapping of cold-induced sweetening in potato using historical phenotypic data. Annals of Applied Biology, 158(3), 248–256. doi:https://doi.org/10.1111/j.1744-7348.2011.00459.x

Biasini, M., Bienert, S., Waterhouse, A., Arnold, K., Studer, G., Schmidt, T.,… Schwede, T. (2014). SWISS-MODEL: modelling protein tertiary and quaternary structure using evolutionary information. Nucleic Acids Research, W252–W258.

Bradburn, S. (2018, November 2). How To Calculate PCR Primer Efficiencies. Retrieved from Top Tip Bio: https://toptipbio.com/calculate-primer-efficiencies/

Braun, D. M., Wang, L., & Ruan, Y.-L. (2014). Understanding and manipulating sucrose phloem loading, unloading, metabolism, and signalling to enhance crop yield and food security. Journal of Experimental Botany, 65(7), 1713–1735. doi:10.1093/jxb/ert416

Brummell, D. A., Chen, R. K., Harris, J. C., Zhang, H., Hamiaux, C., Kralicek, A. V., & McKenzie, M. J. (2011). Induction of vacuolar invertase inhibitor mRNA in potato tubers contributes to cold-induced sweetening resistance and includes spliced hybrid mRNA variants. Journal of Experimental Botany, 62(10), 3519–3534. doi:10.1093/jxb/err043

Castrillon-Arbelaez, P. A., & Delano-Frier, J. P. (2011). The sweet side of inhibition: Invertase inhibitors and their importance in plant development and stress responses. Current Enzyme Inhibition, 7, 169–177.

Chandra, A., Jain, R., & Solomon, S. (2012). Complexities of invertases controlling sucrose accumulation in and retention in sugarcane. Current Science, 102(6), 857–866.

Chourey, P. S., Jain, M., Li, Q. B., & Carlson, S. J. (2006). Genetic control of cell wall invertases in developing endosperm of maize. Planta, 159–167. Retrieved from https://link.springer.com/article/10.1007%2Fs00425-005-0039-5

Datir, S. S., Latimer, J. M., Thompson, S. J., Ridgeway, H. J., Conner, A. J., & Jacobs, J. M. (2012). Allele diversity for the apoplastic invertase inhibitor gene from potato. Molecular Genetics and Genomics, 451–460.

Datir, S. S., Mirikar, D., & RaviKumar, A. (2019). Sequence diversity and in silico structure prediction of the vacuolar invertse inhibitor gene from potato (Solanum tuberosum L.) cultivars differening in sugar content. Food Chemistry, 295, 403–411.

Datir, S., & Ghosh, P. (2020). In silico analysis of the structural diversity and interactions between invertases and invertase inhibitors from potato (Solanum tuberosum L). 3 Biotech, 10(178). doi:https://doi.org/10.1007/s13205-020-02171-y

Emanuelsson, O., Nielsen, H., Brunak, S., & Heijne, G. (2000). Predicting subcellular localization of proteins based on their N-terminal amino acid sequence. Journal of Molecular Biology, 1005–1016.

Gold Biotechnology. (2016, 4 22). Blue-White Screening of Bacterial Colonies Utilizing X-Gal and IPTG Plates. St. Louis, MO, USA. Retrieved from https://www.goldbio.com/documents/1031/Blue%20White%20Screening%20of%20Bacterial%20Colonies%20using%20X-Gal%20and%20IPTG%20Plates.pdf

Greiner, S., Krausgrill, S., & Rausch, T. (1998). Cloning of tobacco apoplasmic invertase inhibitor-proof of function of recombinant protein and expression analysis during plant development. Plant Physiology, 116, 733–742.

Greinet, S., Rausch, T., Sonnewald, U., & Herbers, K. (1999). Ectopic expression of a tobacco invertase inhibitor homolog prevents cold-induced sweetning of potato tubers. Nature Biotechnology, 17, 708–711.

Harvest to Table. (2019). SWEETPOTATOES:SHORT-SUMMER VARIETIES. Retrieved 10 20, 2019, from https://harvesttotable.com/sweet_potatoes_short-summer_va/

Hothorn, M., Van den Ende, W., Lammens, W., Rybin, V., & Scheffzek, K. (2010). Structural insights into the pH-controlled targeting of plant cell wall invertase by a specific inhibitor protein. Proceedings of the National Academy of Science of the United States of America, 107(40), 17427–17432. doi: https://doi.org/10.1073/pnas.1004481107

Hothorn, M., Wolf, S., Aloy, P., Greiner, S., & Scheffzek, K. (2004). Structural Insights into the Target Specificity of Plant Invertase and Pectin Methylesterase Inhibitory Proteins. The Plant Cell, 3437–3447. doi: https://doi.org/10.1105/tpc.104.025684

Huang, G.-J., Sheu, M.-J., Chang, Y.-S., Chang, H.-Y., Huang, S.-S., & Lin, Y.-H. (2008). Isolation and characterisation of invertase inhibitor from sweet potato storage roots. Journal of the Science of Food and Agriculture., 88, 2615–2621. Retrieved from https://onlinelibrary.wiley.com/doi/full/10.1002/jsfa.3380

Huang, G.-J., Sheu, M.-J., Chang, Y.-S., Lu, T.-L., Chang, H.-Y., Huang, S.-S., & Lin, Y.-H. (2008). Isolation and characterisation of invertase inhibitor from sweet potato storage roots. Journal of the Science of Food and Agriculture, 2615–2621. Retrieved from https://citeseerx.ist.psu.edu/viewdoc/download?doi=10.1.1.914.7223&rep=rep1&type=pdf

Jin, Y., Ni, D.-A., & Ruan, Y.-L. (2009). Posttranslational Elevation of Cell Wall Invertase Activity by Silencing Its Inhibitor in Tomato Delays Leaf Senescence and Increases Seed Weight and Fruit Hexose Level. The Plant Cell, 21(7), 2072–2089. doi:10.1105/tpc.108.063719

Jinek, M., Chylinski, K., Fonfara, I., Hauer, M., Doudna, J. A., & Charpentier, E. (2012). A programmable dual-RNA-guided DNA endonuclease in adaptive bacterial immunity. Science, 816–821. doi:10.1126/science.1225829

Karuri, H. W., Ateka, E. M., Amata, R., Nyende, A. B., Muigai, A. W., Mwasame, E., & Gichuki, S. T. (2010). Evaluating Diversity among Kenyan Sweet Potato Genotypes Using Morphological and SSR Markers. International Journal of Agriculture & Biology, 12, 33–38. Retrieved from http://www.fspublishers.org/published_papers/31561_.pdf

Kelly, L. A., Mezulis, S., Yates, C. M., Wass, M. N., & Sternberg, M. J. (2015). The Phyre2 web portal for protein modeling, prediction and analysis. 845–858.

Kiktev, D. A., Sheng, Z., Lobachev, K. S., & Petes, T. D. (2018). GC content elevates mutation and recombination rates in the yeast Saccharomyces cerevisiae. Proceedings of the National Academy of Sciences, 115(30), E7109–E7118. doi:https://doi.org/10.1073/pnas.1807334115

Kocal, N., Sonnewald, U., & Sonnewald, S. (2008). Cell Wall-Bound Invertase Limits Sucrose Export and Is Involved in Symptom Development and Inhibition of Photosynthesis during Compatible Interaction between Tomato and Xanthomonas campestris pv vesicatoria. Plant Physiology, 148(3), 1523–1536. doi: https://doi.org/10.1104/pp.108.127977

Kofi Anan Foundation. (2021, April 20). Combatting Hunger. Retrieved November 20, 2021, from New project: Promoting Orange-Fleshed Sweet Potato in Ghana: https://www.kofiannanfoundation.org/combatting-hunger/orange-fleshed-sweet-potato-ghana/

Krausgrill, S., Sander, A., Greiner, S., Weil, M., & Rausch, T. (1996). Regulation of cell wall invertase by a proteinaceous inhibitor. Journal of Experimental Botany, 1193–1198. Retrieved from https://academic.oup.com/jxb/article/47/Special_Issue/1193/461050

Lareo, C., & Ferrari, M. D. 2019). Sweet Potato as a Bioenergy Crop for Fuel Ethanol Production: Perspectives and Challenges. In S. Ramachandran, & R. C. Ray, Bioethanol Production from Food Crops Sustainable Sources, Interventions, and Challenges (pp. 115–147). Academic Press. doi:https://doi.org/10.1016/B978-0-12-813766-6.00007-2

Li, M., Yang, S., Xu, W., Pu, Z., Feng, J., Wang, Z.,… Tan, W. (2019). The wild sweetpotato (Ipomoea trifida) genome provides insights into storage root development. BMC Plant Biology, 19, 119. Retrieved from https://bmcplantbiol.biomedcentral.com/articles/10.1186/s12870-019-1708-z

Link, M., Rausch, T., & Greiner, S. (2004). InArabidopsis thaliana, the invertase inhibitors AtC/VIF1 and 2exhibit distinct target enzyme specificities and expression profiles. FEBS Letters, 573, 105–109. doi:10.1016/j.febslet.2004.07.062

LSU AgCenter. (2019). Sweet Potato Variety Descriptions. Retrieved 10 30, 2019, from Sweet Potato Varieties: https://www.lsuagcenter.com/portals/our_offices/research_stations/sweetpotato/features/varieties/sweet-potato-variety-descriptions

Madeira, F., Park, Y. M., Lee, J., Buso, N., Gur, T., Madhusoodanan, N.,… Lopez, R. (2019). The EMBL-EBI search and sequence analysis tools APIs in 2019. Nucleic Acids Research. doi:10.1093/nar/gkz268

Michigan State University. (2016, August 1). Sweetpotato Genomics Resource at Michigan State University. Retrieved from Sweetpotato Genomics Resource: http://sweetpotato.uga.edu/

Moghaddam, M. R., & Ende, W. V. (2012). Sugars and plant innate immunity. Journal of Experimental Botany, 63(11), 3989–3998. doi:10.1093/jxb/ers129

North Carolina State University. (2018, Winter). Unburried treasure:Breeding better sweet potatoes for the world. NC STATE Alumni Magazine. Retrieved from https://www.sweetpotatoknowledge.org/wp-content/uploads/2019/02/Breeding-better.pdf

Paliwal, P., Jain, D., Joshi, A., Ameta, K. D., Chaudhary, R., & Singh, A. (2020). Diversity analysis of Sweet Potato (Ipomoea batatas [L.] Lam) genotypes using morphological, biochemical and molecular markers. Indian Journal of Experimental Biology, 58, 276–285. Retrieved from http://nopr.niscair.res.in/bitstream/123456789/54257/1/IJEB%2058%20%284%29%20276-285.pdf

Pope, D. T., Nielsen, L. W., & Miller, N. C. (1971). Jewel, a new sweet potato variety for North Carolina. 442(11).

Pressey, R. (1967). Invertase inhibitor from potatoes:purification, characterization, and reactivity with plant invertases. Plant Physiology, 42, 1780–1786.

Qin, G., Zhu, Z., Wang, Z., Cai, J., Chen, Y., Li, L., & Tian, S. (2016). A Tomato Vacuolar Invertase Inhibitor Mediates Sucrose Metabolismand Influences Fruit Ripening. Plant Physiology, 172, 1596–1611. Retrieved from https://www.plantphysiol.org

Rao, Y. S., Chai, X. W., Wang, Z. F., Nie, H. Q., & Zhang, X. Q. (2013). Impact of GC content on gene expression pattern in chicken. Genetics Selection Evolution, 45(9). Retrieved from http://www.gsejournal.org/content/45/1/9

Rastogi, S., & Liberles, D. A. (2005). Subfunctionalization of duplicated genes as a transition state to neofunctionalization. BMC Evolutionary Biology, 5(28). doi:10.1186/1471-2148-5-28

Rausch, T., & Greiner, S. (2004). Plant protein inhibitors of invertases. Biochimica et Biophysica Acta - Proteins and Proteomics, 1696(2), 253–261. doi:https://doi.org/10.1016/j.bbapap.2003.09.017

Rolston, L. H., Clark, C. A., Cannon, J. M., Randle, W. M., Riley, E. G., Wilson, P. W., & Robbins, M. L. (1987). ‘Beauregard’ Sweetpotato. HortScience, 22, 1338–1339.

Ruan, Y. L., Jin, Y., Yang, Y.-J., Li, G.-J., & Boyer, J. S. (2010). Sugar input, metabolism, and signaling mediated by invertase: roles in development, yield potential, and response to drought and heat. Molecular Plant, 3, 942–955.

Sander, A., Krausgrill, S., Greiner, S., Weil, M., & Rausch, T. (1996, May). Sucrose protects cell wall invertase but not vacuolar invertase against proteinaceous inhibitors. 385(3), 171–175. doi:10.1016/0014-5793(96)00378-x

Scripps Networks, LLC. (2019). Top 5 Orange SweetPotatoes to Grow. Retrieved 10 30, 2019, from Top 5 Orange SweetPotatoes to Grow: https://www.diynetwork.com/how-to/outdoors/gardening/top-5-orange-sweet-potatoes-to-grow

Shepherd, L. V., Bradshaw, J. E., Dale, M. F., McNicol, J. W., Pont, S. D., Mottram, D. S., & Davies, H. V. (2010). Variation in acrylamide producing potential in potato: Segregation of the trait in a breeding population. Food Chemistry, 123(3), 568–573. doi:10.1016/j.foodchem.2010.04.070

Shivalingamurthy, S. G., Anangi, R., Kalaipandian, S., Glassop, D., King, G. F., & Rae, A. L. (2018). Identification and Functional Characterization of Sugarcane Invertase Inhibitor (ShINH1): A Potential Candidate for Reducing Pre-and Post-harvest Loss of Sucrose in Sugarcane. Frontiers in Plant Science, 9. doi:10.3389/fpls.2018.00598

Smarda, P., Bures, P., Horova, L., Leitch, l. J., Mucina, L., Pacini, E.,… Rotreklova, O. (2014). Ecological and evolutionary significance of genomic GC content diversity in monocots. PNAS. doi:10.1073/pnas.1321152111

Solomon, S. (2009). Post-harvest deterioration in sugarcane. Sugar Technology, 11(2), 109–123.

Song, H., Gao, H., Liu, J., Tian, P., & Nan, Z. (2017). Comprehensive analysis of correlations among codon usage bias, gene expression,and substitution rate in Arachis duranensis and Arachis ipaёnsis orthologs. Nature Scientifc Report. doi:10.1038/s41598-017-13981-1

Sturm, A. (1999). Invertases: Primary structures, functions, and roles in plant development andsucrose partitioning. Plant Physiology, 121(1), 1–8. Retrieved from https://www.ncbi.nlm.nih.gov/pubmed/10482654

Su, T., Han, M., Min, J., Zhou, H., Zhang, Q., Zhao, J., & Fang, Y. (2020). Functional Characterization of Invertase Inhibitors PtC/VIF1 and 2 Revealed Their Involvements in the Defense Response to Fungal Pathogen in Populus trichocarpa. Frontiers in Plant Science. doi:https://doi.org/10.3389/fpls.2019.01654

Tauzin, A. S., & Giardina, T. (2014). Sucrose and invertases, a part of the plant defense response to the biotic stresses. Frontiers in Plant Sciences. doi:https://doi.org/10.3389/fpls.2014.00293

Tauzin, A. S., & Giardina, T. (2014). Sucrose and invertases, a part of the plant defenseresponse to the biotic stresses. Frontiers in Plant Science, 5, 293. Retrieved from https://www.ncbi.nlm.nih.gov/pmc/articles/PMC4066202/

Veillet, F., Gaillard, C., Coutos-Thévenot, P., & Camera, S. L. (2016). Targeting the AtCWIN1 Gene to Explore the Role of Invertases in Sucrose Transport in Roots and during Botrytis cinerea Infection. Frontiers in Plant Science. doi:https://doi.org/10.3389/fpls.2016.01899

Wu, S., Lau, K. H., Cao, Q., Hamilton, J. P., Sun, H., Zhou, C.,… Fei, Z. (2018). Genome sequences of two diploid wild relatives of cultivated sweetpotato reveal targets for genetic improvement. Nature Communications, 9(4580). Retrieved from https://www.nature.com/articles/s41467-018-06983-8

Xu, X.-x., Hu, Q., Yang, W.-n., & Jin, Y. (2017). The roles of cell wall invertase inhibitor in regulating chilling tolerance in tomato. BMC Plant Biology. doi:10.1186/s12870-017-1145-9

Yang, J., Moeinzadeh, M.-H., Kuhl, H., Helmuth, J., Xiao, P., Haas, S.,… Timmermann, B. (2017). Haplotype-resolved sweet potato genome traces back its hexaploidization history. Nature Plants. doi:10.1038/s41477-017-0002-z

Zhang, N., Jiang, J., Yang, Y.-l., & Wang, Z.-h. (2015). Functional characterization of an invertase inhibitor gene involved in sucrose metabolism in tomato fruit. Journal of Zhejiang University Science B, 845–856. doi:10.1631/jzus.B1400319

Zrenner, R., Schuler, K., & Sonnewald, U. (1996). Soluble acid inveratse determines the hexose-to-sucrose ratio in cold-stored potato tubers. Planta, 258, 287–292.

