## Supplementary File S1 for "Diversity, Phylogenetic Relationships, And Expression Profiles Of Invertase Inhibitor Genes In Sweetpotato"

DNA sequence alignment of invertase inhibitor genes in sweetpotatoes. Sequences from other species were added as outgroups. **Accession numbers are preceded by 2 or 3-letters to identify the species. Iba, *Ipomoea batatas*; Itf, *Ipomoea trifida*; Itb, *Ipomoea triloba*; Sl, *Solanum lycopersicum*; Ath, *Arabidopsis thaliana***

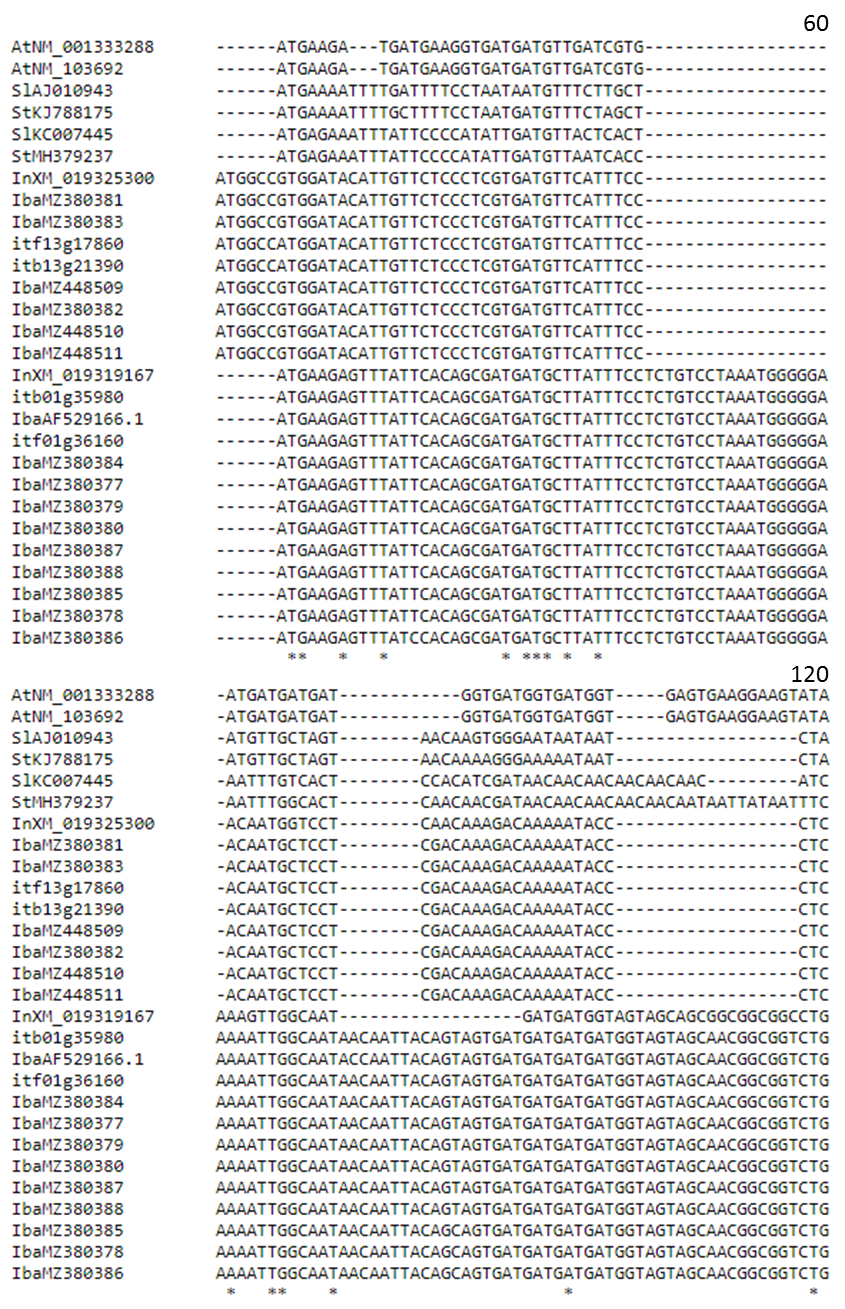

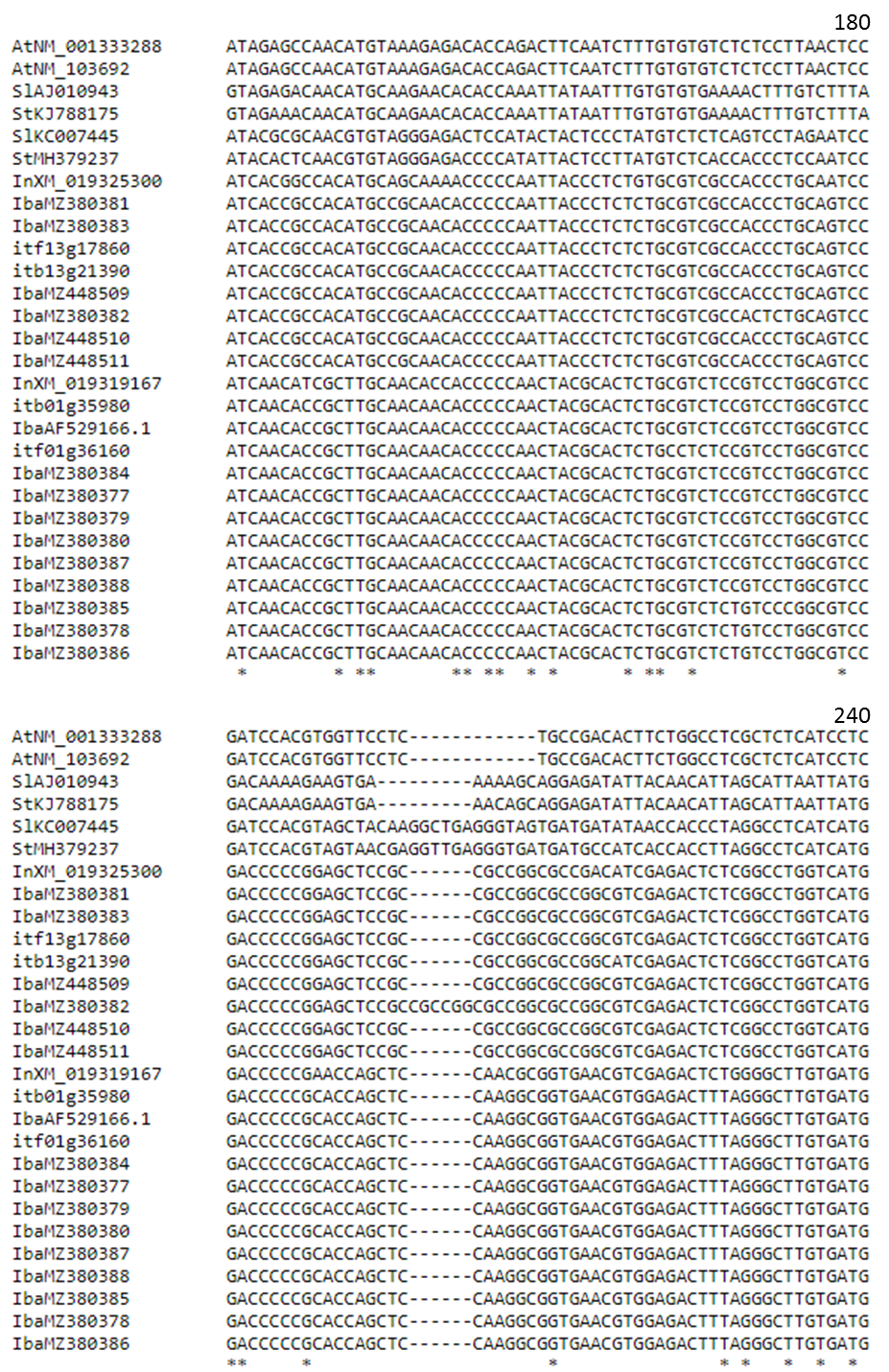

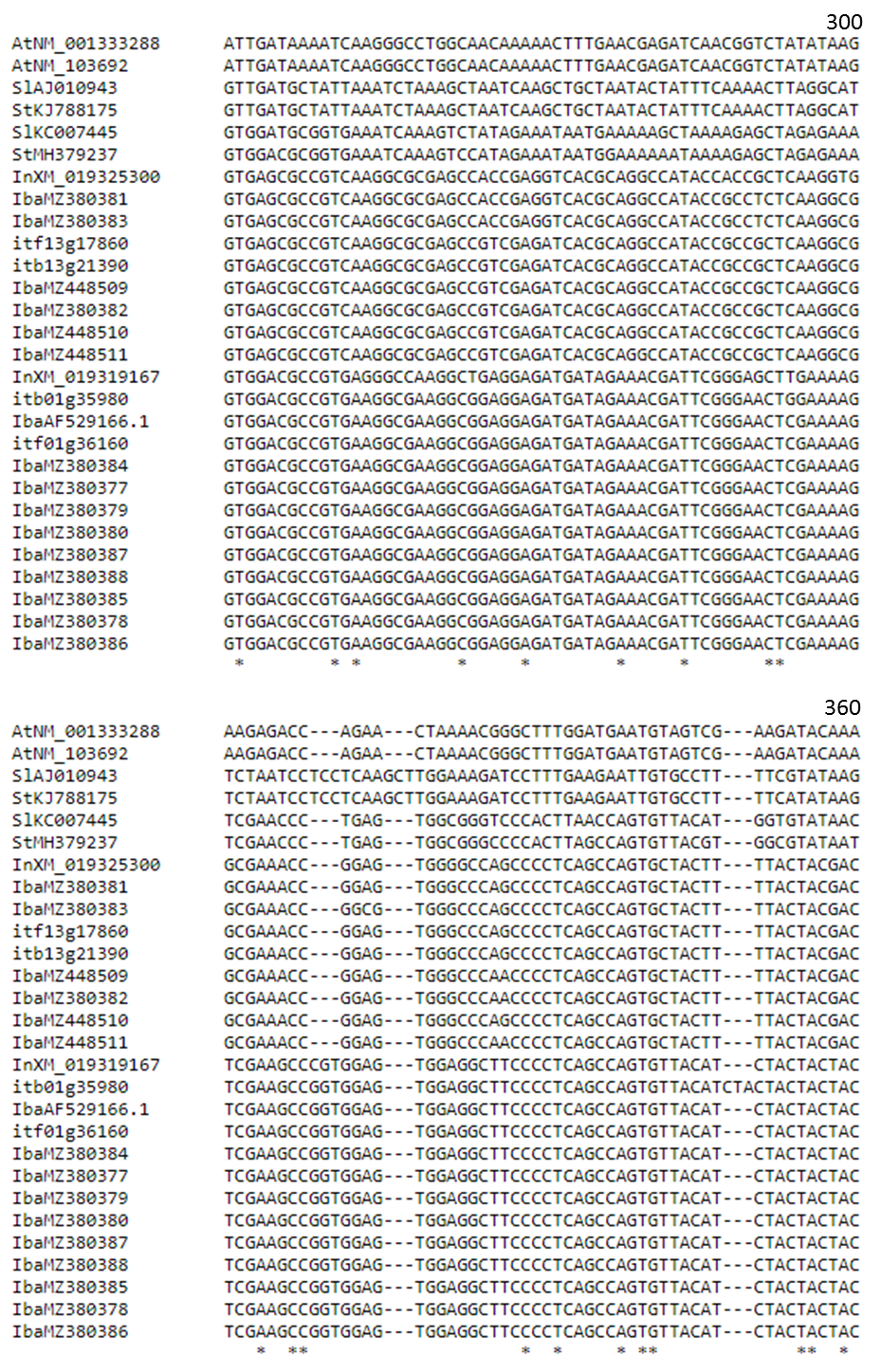

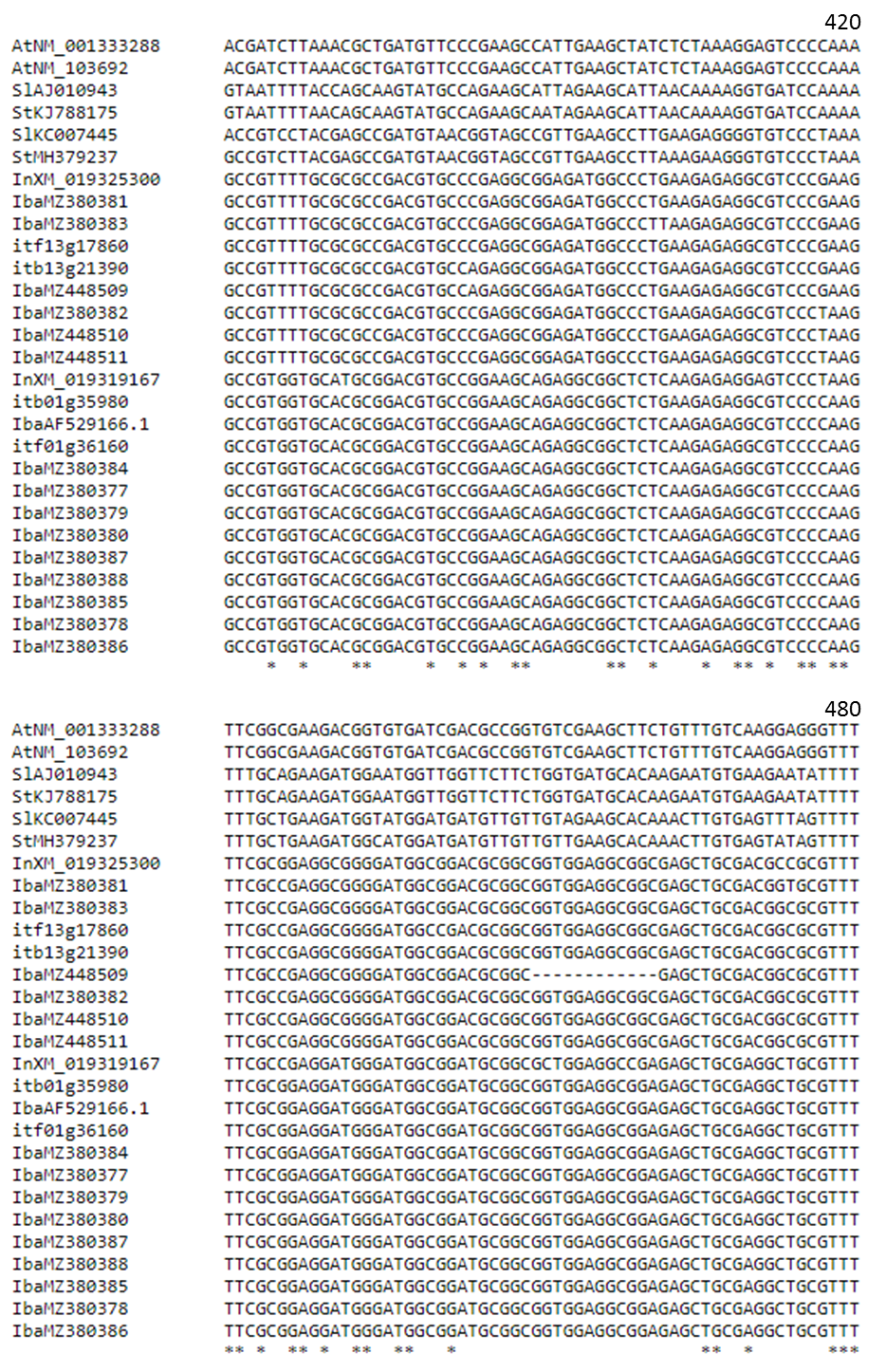

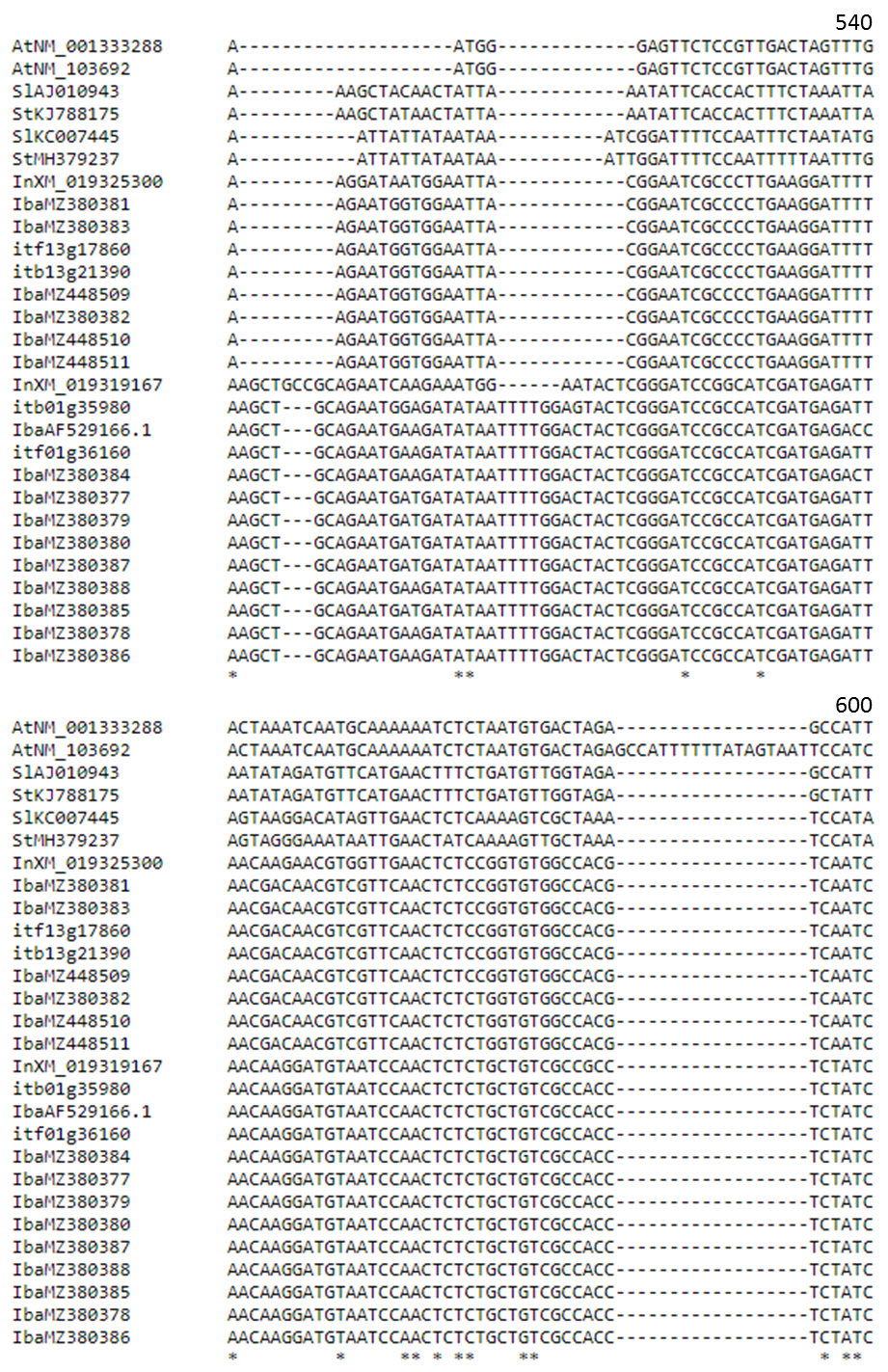

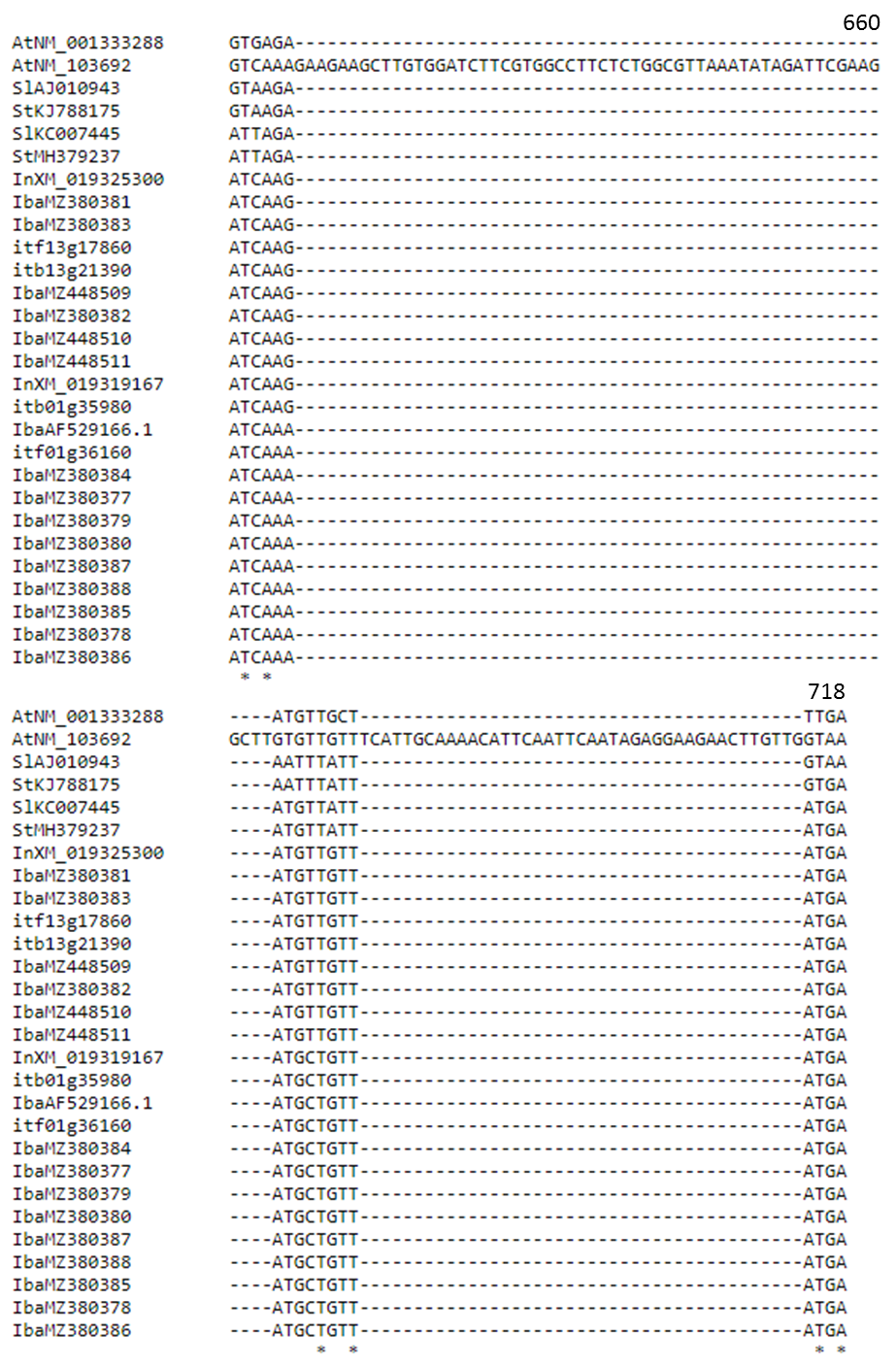
