## Supplementary File S2 for "Diversity, Phylogenetic Relationships, And Expression Profiles Of Invertase Inhibitor Genes In Sweetpotato"

### SPITI1

```
MZ380381_J10      CAAAGATGGCCGTGGATACATTGTTCTCCCTCGTGATGTTCATTTCCACAATGCTCCTCG
MZ380383_J4       CAAAGATGGCCGTGGATACATTGTTCTCCCTCGTGATGTTCATTTCCACAATGCTCCTCG
MZ448509_B3       CAAAGATGGCCGTGGATACATTGTTCTCCCTCGTGATGTTCATTTCCACAATGCTCCTCG
MZ448510_B2       CAAAGATGGCCGTGGATACATTGTTCTCCCTCGTGATGTTCATTTCCACAATGCTCCTCG
MZ380382_J9       CAAAGATGGCCGTGGATACATTGTTCTCCCTCGTGATGTTCATTTCCACAATGCTCCTCG
MZ448511_B1       CAAAGATGGCCGTGGATACATTGTTCTCCCTCGTGATGTTCATTTCCACAATGCTCCTCG
                  ************************************************************

MZ380381_J10      ACAAAGACAAAAATACCCTCATCACCGCCACATGCCGCAACACCCCCAATTACCCTCTCT
MZ380383_J4       ACAAAGACAAAAATACCCTCATCACCGCCACATGCCGCAACACCCCCAATTACCCTCTCT
MZ448509_B3       ACAAAGACAAAAATACCCTCATCACCGCCACATGCCGCAACACCCCCAATTACCCTCTCT
MZ448510_B2       ACAAAGACAAAAATACCCTCATCACCGCCACATGCCGCAACACCCCCAATTACCCTCTCT
MZ380382_J9       ACAAAGACAAAAATACCCTCATCACCGCCACATGCCGCAACACCCCCAATTACCCTCTCT
MZ448511_B1       ACAAAGACAAAAATACCCTCATCACCGCCACATGCCGCAACACCCCCAATTACCCTCTCT
                  ************************************************************

MZ380381_J10      GCGTCGCCACCCTGCAGTCCGACCCCCGGAGCTCCGC------CGCCGGCGCCGGCGTCG
MZ380383_J4       GCGTCGCCACCCTGCAGTCCGACCCCCGGAGCTCCGC------CGCCGGCGCCGGCGTCG
MZ448509_B3       GCGTCGCCACCCTGCAGTCCGACCCCCGGAGCTCCGC------CGCCGGCGCCGGCGTCG
MZ448510_B2       GCGTCGCCACCCTGCAGTCCGACCCCCGGAGCTCCGC------CGCCGGCGCCGGCGTCG
MZ380382_J9       GCGTCGCCACTCTGCAGTCCGACCCCCGGAGCTCCGCCGCCGGCGCCGGCGCCGGCGTCG
MZ448511_B1       GCGTCGCCACCCTGCAGTCCGACCCCCGGAGCTCCGC------CGCCGGCGCCGGCGTCG
                  ********** **************************      *****************

MZ380381_J10      AGACTCTCGGCCTGGTCATGGTGAGCGCCGTCAAGGCGCGAGCCACCGAGGTCACGCAGG
MZ380383_J4       AGACTCTCGGCCTGGTCATGGTGAGCGCCGTCAAGGCGCGAGCCACCGAGGTCACGCAGG
MZ448509_B3       AGACTCTCGGCCTGGTCATGGTGAGCGCCGTCAAGGCGCGAGCCGTCGAGATCACGCAGG
MZ448510_B2       AGACTCTCGGCCTGGTCATGGTGAGCGCCGTCAAGGCGCGAGCCGTCGAGATCACGCAGG
MZ380382_J9       AGACTCTCGGCCTGGTCATGGTGAGCGCCGTCAAGGCGCGAGCCGTCGAGATCACGCAGG
MZ448511_B1       AGACTCTCGGCCTGGTCATGGTGAGCGCCGTCAAGGCGCGAGCCGTCGAGATCACGCAGG
                  ********************************************  **** *********

MZ380381_J10      CCATACCGCCTCTCAAGGCGGCGAAACCGGAGTGGGCCCAGCCCCTCAGCCAGTGCTACT
MZ380383_J4       CCATACCGCCTCTCAAGGCGGCGAAACCGGCGTGGGCCCAGCCCCTCAGCCAGTGCTACT
MZ448509_B3       CCATACCGCCGCTCAAGGCGGCGAAACCGGAGTGGGCCCAACCCCTCAGCCAGTGCTACT
MZ448510_B2       CCATACCGCCGCTCAAGGCGGCGAAACCGGAGTGGGCCCAGCCCCTCAGCCAGTGCTACT
MZ380382_J9       CCATACCGCCGCTCAAGGCGGCGAAACCGGAGTGGGCCCAACCCCTCAGCCAGTGCTACT
MZ448511_B1       CCATACCGCCGCTCAAGGCGGCGAAACCGGAGTGGGCCCAACCCCTCAGCCAGTGCTACT
                  ********** ******************* ********* *******************

MZ380381_J10      TTTACTACGACGCCGTTTTGCGCGCCGACGTGCCCGAGGCGGAGATGGCCCTGAAGAGAG
MZ380383_J4       TTTACTACGACGCCGTTTTGCGCGCCGACGTGCCCGAGGCGGAGATGGCCCTTAAGAGAG
MZ448509_B3       TTTACTACGACGCCGTTTTGCGCGCCGACGTGCCAGAGGCGGAGATGGCCCTGAAGAGAG
MZ448510_B2       TTTACTACGACGCCGTTTTGCGCGCCGACGTGCCCGAGGCGGAGATGGCCCTGAAGAGAG
MZ380382_J9       TTTACTACGACGCCGTTTTGCGCGCCGACGTGCCCGAGGCGGAGATGGCCCTGAAGAGAG
MZ448511_B1       TTTACTACGACGCCGTTTTGCGCGCCGACGTGCCCGAGGCGGAGATGGCCCTGAAGAGAG
                  ********************************** ***************** *******

MZ380381_J10      GCGTCCCGAAGTTCGCCGAGGCGGGGATGGCGGACGCGGCGGTGGAGGCGGCGAGCTGCG
MZ380383_J4       GCGTCCCGAAGTTCGCCGAGGCGGGGATGGCGGACGCGGCGGTGGAGGCGGCGAGCTGCG
MZ448509_B3       GCGTCCCGAAGTTCGCCGAGGCGGGGATGGCGGAC------------GCGGCGAGCTGCG
MZ448510_B2       GCGTCCCTAAGTTCGCCGAGGCGGGGATGGCGGACGCGGCGGTGGAGGCGGCGAGCTGCG
MZ380382_J9       GCGTCCCTAAGTTCGCCGAGGCGGGGATGGCGGACGCGGCGGTGGAGGCGGCGAGCTGCG
MZ448511_B1       GCGTCCCTAAGTTCGCCGAGGCGGGGATGGCGGACGCGGCGGTGGAGGCGGCGAGCTGCG
                  ******* ***************************            *************

MZ380381_J10      ACGGTGCGTTTAAGAATGGTGGAATTACGGAATCGCCCCTGAAGGATTTTAACGACAACG
MZ380383_J4       ACGGCGCGTTTAAGAATGGTGGAATTACGGAATCGCCCCTGAAGGATTTTAACGACAACG
MZ448509_B3       ACGGCGCGTTTAAGAATGGTGGAATTACGGAATCGCCCCTGAAGGATTTTAACGACAACG
MZ448510_B2       ACGGCGCGTTTAAGAATGGTGGAATTACGGAATCGCCCCTGAAGGATTTTAACGACAACG
MZ380382_J9       ACGGCGCGTTTAAGAATGGTGGAATTACGGAATCGCCCCTGAAGGATTTTAACGACAACG
MZ448511_B1       ACGGCGCGTTTAAGAATGGTGGAATTACGGAATCGCCCCTGAAGGATTTTAACGACAACG
                  **** *******************************************************


MZ380381_J10      TCGTTCAACTCTCCGGTGTGGCCACGTCAATCATCAAGATGTTGTTATGA
MZ380383_J4       TCGTTCAACTCTCCGGTGTGGCCACGTCAATCATCAAGATGTTGTTATGA
MZ448509_B3       TCGTTCAACTCTCCGGTGTGGCCACGTCAATCATCAAGATGTTGTTATGA
MZ448510_B2       TCGTTCAACTCTCTGGTGTGGCCACGTCAATCATCAAGATGTTGTTATGA
MZ380382_J9       TCGTTCAACTCTCTGGTGTGGCCACGTCAATCATCAAGATGTTGTTATGA
MZ448511_B1       TCGTTCAACTCTCTGGTGTGGCCACGTCAATCATCAAGATGTTGTTATGA
                  ************* ************************************
```

### SPITI2

```
P20_J8          GGATCGGCTCTCATCGATCTGATCTCTA-TCTACACCAAATTACTGC-----AAGATGAA
P20_J1          GGATCGGCTCTCATCGATCTGATCTTTAATTTATACCAAATTACTGCAAGCTAAGATGAA
P20_B9          GGATCGGCTCTCATCGATCTGATCTTTAATTTACACCAAATTACTGCAAGCTAAGATGAA
P20_B2          GGATCGGCTCTCATCGATCTGATCTTTAATTTACACCAAATTACTGGAAGCTAAGATGAA
P20_J3          GGATCGGCTCTCATCGATCTGATCTTTAATTTACACCAAATTACTGGAAGCTAAGATGAA
P20_B1          GGATCGGCTCTCATCGATCTGGTCTTTAATTTACACCAAATTACTGGAAGCTAAGATGAA
P20_J7          GGATCGGCTCTCATCGATCTGATC-----TCTACACCAAATTACTGC-----AAGATGAA
P20_B3          GGATCGGCTCTCATCGATCTGATC-----TCTACACCAAATTACTGC-----AAGATGAA
P20_J5          GGATCGGCTCTCATCGATCTGATC-----TCTACACCAAATTACTGC-----AAGATGAA
                ********************* **     * ** ************      ********

P20_J8          GAGTTTATTCACAGCGATGATGCTTATTTCCTCTGTCCTAAATGGGGGAAAAATTGGCAA
P20_J1          GAGTTTATTCACAGCGATGATGCTTATTTCCTCTGTCCTAAATGGGGGAAAAATTGGCAA
P20_B9          GAGTTTATTCACAGCGATGATGCTTATTTCCTCTGTCCTAAATGGGGGAAAAATTGGCAA
P20_B2          GAGTTTATTCACAGCGATGATGCTTATTTCCTCTGTCCTAAATGGGGGAAAAATTGGCAA
P20_J3          GAGTTTATTCACAGCGATGATGCTTATTTCCTCTGTCCTAAATGGGGGAAAAATTGGCAA
P20_B1          GAGTTTATTCACAGCGATGATGCTTATTTCCTCTGTCCTAAATGGGGGAAAAATTGGCAA
P20_J7          GAGTTTATTCACAGCGATGATGCTTATTTCCTCTGTCCTAAATGGGGGAAAAATTGGCAA
P20_B3          GAGTTTATTCACAGCGATGATGCTTATTTCCTCTGTCCTAAATGGGGGAAAAATTGGCAA
P20_J5          GAGTTTATCCACAGCGATGATGCTTATTTCCTCTGTCCTAAATGGGGGAAAAATTGGCAA
                ******** ***************************************************

P20_J8          TAACAATTACAGTAGTGATGATGATGATGGTAGTAGCAACGGCGGTCTGATCAACACCGC
P20_J1          TAACAATTACAGTAGTGATGATGATGATGGTAGTAGCAACGGCGGTCTGATCAACACCGC
P20_B9          TAACAATTACAGTAGTGATGATGATGATGGTAGTAGCAACGGCGGTCTGATCAACACCGC
P20_B2          TAACAATTACAGTAGTGATGATGATGATGGTAGTAGCAACGGCGGTCTGATCAACACCGC
P20_J3          TAACAATTACAGTAGTGATGATGATGATGGTAGTAGCAACGGCGGTCTGATCAACACCGC
P20_B1          TAACAATTACAGTAGTGATGATGATGATGGTAGTAGCAACGGCGGTCTGATCAACACCGC
P20_J7          TAACAATTACAGCAGTGATGATGATGATGGTAGTAGCAACGGCGGTCTGATCAACACCGC
P20_B3          TAACAATTACAGCAGTGATGATGATGATGGTAGTAGCAACGGCGGTCTGATCAACACCGC
P20_J5          TAACAATTACAGCAGTGATGATGATGATGGTAGTAGCAACGGCGGTCTGATCAACACCGC
                ************ ***********************************************

P20_J8          TTGCAACAACACCCCCAACTACGCACTCTGCGTCTCCGTCCTGGCGTCCGACCCCCGCAC
P20_J1          TTGCAACAACACCCCCAACTACGCACTCTGCGTCTCCGTCCTGGCGTCCGACCCCCGCAC
P20_B9          TTGCAACAACACCCCCAACTACGCACTCTGCGTCTCCGTCCTGGCGTCCGACCCCCGCAC
P20_B2          TTGCAACAACACCCCCAACTACGCACTCTGCGTCTCCGTCCTGGCGTCCGACCCCCGCAC
P20_J3          TTGCAACAACACCCCCAACTACGCACTCTGCGTCTCCGTCCTGGCGTCCGACCCCCGCAC
P20_B1          TTGCAACAACACCCCCAACTACGCACTCTGCGTCTCCGTCCTGGCGTCCGACCCCCGCAC
P20_J7          TTGCAACAACACCCCCAACTACGCACTCTGCGTCTCTGTCCCGGCGTCCGACCCCCGCAC
P20_B3          TTGCAACAACACCCCCAACTACGCACTCTGCGTCTCTGTCCTGGCGTCCGACCCCCGCAC
P20_J5          TTGCAACAACACCCCCAACTACGCACTCTGCGTCTCTGTCCTGGCGTCCGACCCCCGCAC
                ************************************ **** ******************

P20_J8          CAGCTCCAAGGCGGTGAACGTGGAGACTTTAGGGCTTGTGATGGTGGACGCCGTGAAGGC
P20_J1          CAGCTCCAAGGCGGTGAACGTGGAGACTTTAGGGCTTGTGATGGTGGACGCCGTGAAGGC
P20_B9          CAGCTCCAAGGCGGTGAACGTGGAGACTTTAGGGCTTGTGATGGTGGACGCCGTGAAGGC
P20_B2          CAGCTCCAAGGCGGTGAACGTGGAGACTTTAGGGCTTGTGATGGTGGACGCCGTGAAGGC
P20_J3          CAGCTCCAAGGCGGTGAACGTGGAGACTTTAGGGCTTGTGATGGTGGACGCCGTGAAGGC
P20_B1          CAGCTCCAAGGCGGTGAACGTGGAGACTTTAGGGCTTGTGATGGTGGACGCCGTGAAGGC
P20_J7          CAGCTCCAAGGCGGTGAACGTGGAGACTTTAGGGCTTGTGATGGTGGACGCCGTGAAGGC
P20_B3          CAGCTCCAAGGCGGTGAACGTGGAGACTTTAGGGCTTGTGATGGTGGACGCCGTGAAGGC
P20_J5          CAGCTCCAAGGCGGTGAACGTGGAGACTTTAGGGCTTGTGATGGTGGACGCCGTGAAGGC
                ************************************************************

P20_J8          GAAGGCGGAGGAGATGATAGAAACGATTCGGGAACTCGAAAAGTCGAAGCCGGTGGAGTG
P20_J1          GAAGGCGGAGGAGATGATAGAAACGATTCGGGAACTCGAAAAGTCGAAGCCGGTGGAGTG
P20_B9          GAAGGCGGAGGAGATGATAGAAACGATTCGGGAACTCGAAAAGTCGAAGCCGGTGGAGTG
P20_B2          GAAGGCGGAGGAGATGATAGAAACGATTCGGGAACTCGAAAAGTCGAAGCCGGTGGAGTG
P20_J3          GAAGGCGGAGGAGATGATAGAAACGATTCGGGAACTCGAAAAGTCGAAGCCGGTGGAGTG
P20_B1          GAAGGCGGAGGAGATGATAGAAACGATTCGGGAACTCGAAAAGTCGAAGCCGGTGGAGTG
P20_J7          GAAGGCGGAGGAGATGATAGAAACGATTCGGGAACTCGAAAAGTCGAAGCCGGTGGAGTG
P20_B3          GAAGGCGGAGGAGATGATAGAAACGATTCGGGAACTCGAAAAGTCGAAGCCGGTGGAGTG
P20_J5          GAAGGCGGAGGAGATGATAGAAACGATTCGGGAACTCGAAAAGTCGAAGCCGGTGGAGTG
                ************************************************************


P20_J8          GAGGCTTCCCCTCAGCCAGTGTTACATCTACTACTACGCCGTGGTGCACGCGGACGTGCC
P20_J1          GAGGCTTCCCCTCAGCCAGTGTTACATCTACTACTACGCCGTGGTGCACGCGGACGTGCC
P20_B9          GAGGCTTCCCCTCAGCCAGTGTTACATCTACTACTACGCCGTGGTGCACGCGGACGTGCC
P20_B2          GAGGCTTCCCCTCAGCCAGTGTTACATCTACTACTACGCCGTGGTGCACGCGGACGTGCC
P20_J3          GAGGCTTCCCCTCAGCCAGTGTTACATCTACTACTACGCCGTGGTGCACGCGGACGTGCC
P20_B1          GAGGCTTCCCCTCAGCCAGTGTTACATCTACTACTACGCCGTGGTGCACGCGGACGTGCC
P20_J7          GAGGCTTCCCCTCAGCCAGTGTTACATCTACTACTACGCCGTGGTGCACGCGGACGTGCC
P20_B3          GAGGCTTCCCCTCAGCCAGTGTTACATCTACTACTACGCCGTGGTGCACGCGGACGTGCC
P20_J5          GAGGCTTCCCCTCAGCCAGTGTTACATCTACTACTACGCCGTGGTGCACGCGGACGTGCC
                ************************************************************

P20_J8          GGAAGCAGAGGCGGCTCTCAAGAGAGGCGTCCCCAAGTTCGCGGAGGATGGGATGGCGGA
P20_J1          GGAAGCAGAGGCGGCTCTCAAGAGAGGCGTCCCCAAGTTCGCGGAGGATGGGATGGCGGA
P20_B9          GGAAGCAGAGGCGGCTCTCAAGAGAGGCGTCCCCAAGTTCGCGGAGGATGGGATGGCGGA
P20_B2          GGAAGCAGAGGCGGCTCTCAAGAGAGGCGTCCCCAAGTTCGCGGAGGATGGGATGGCGGA
P20_J3          GGAAGCAGAGGCGGCTCTCAAGAGAGGCGTCCCCAAGTTCGCGGAGGATGGGATGGCGGA
P20_B1          GGAAGCAGAGGCGGCTCTCAAGAGAGGCGTCCCCAAGTTCGCGGAGGATGGGATGGCGGA
P20_J7          GGAAGCAGAGGCGGCTCTCAAGAGAGGCGTCCCCAAGTTCGCGGAGGATGGGATGGCGGA
P20_B3          GGAAGCAGAGGCGGCTCTCAAGAGAGGCGTCCCCAAGTTCGCGGAGGATGGGATGGCGGA
P20_J5          GGAAGCAGAGGCGGCTCTCAAGAGAGGCGTCCCCAAGTTCGCGGAGGATGGGATGGCGGA
                ************************************************************

P20_J8          TGCGGCGGTGGAGGCGGAGAGCTGCGAGGCTGCGTTTAAGCTGCAGAATGAAGATATAAT
P20_J1          TGCGGCGGTGGAGGCGGAGAGCTGCGAGGCTGCGTTTAAGCTGCAGAATGAAGATATAAT
P20_B9          TGCGGCGGTGGAGGCGGAGAGCTGCGAGGCTGCGTTTAAGCTGCAGAATGATGATATAAT
P20_B2          TGCGGCGGTGGAGGCGGAGAGCTGCGAGGCTGCGTTTAAGCTGCAGAATGATGATATAAT
P20_J3          TGCGGCGGTGGAGGCGGAGAGCTGCGAGGCTGCGTTTAAGCTGCAGAATGATGATATAAT
P20_B1          TGCGGCGGTGGAGGCGGAGAGCTGCGAGGCTGCGTTTAAGCTGCAGAATGATGATATAAT
P20_J7          TGCGGCGGTGGAGGCGGAGAGCTGCGAGGCTGCGTTTAAGCTGCAGAATGATGATATAAT
P20_B3          TGCGGCGGTGGAGGCGGAGAGCTGCGAGGCTGCGTTTAAGCTGCAGAATGAAGATATAAT
P20_J5          TGCGGCGGTGGAGGCGGAGAGCTGCGAGGCTGCGTTTAAGCTGCAGAATGAAGATATAAT
                *************************************************** ********

P20_J8          TTTGGACTACTCGGGATCCGCCATCGATGAGACTAACAAGGATGTAATCCAACTCTCTGC
P20_J1          TTTGGACTACTCGGGATCCGCCATCGATGAGATTAACAAGGATGTAATCCAACTCTCTGC
P20_B9          TTTGGACTACTCGGGATCCGCCATCGATGAGATTAACAAGGATGTAATCCAACTCTCTGC
P20_B2          TTTGGACTACTCGGGATCCGCCATCGATGAGATTAACAAGGATGTAATCCAACTCTCTGC
P20_J3          TTTGGACTACTCGGGATCCGCCATCGATGAGATTAACAAGGATGTAATCCAACTCTCTGC
P20_B1          TTTGGACTACTCGGGATCCGCCATCGATGAGATTAACAAGGATGTAATCCAACTCTCTGC
P20_J7          TTTGGACTACTCGGGATCCGCCATCGATGAGATTAACAAGGATGTAATCCAACTCTCTGC
P20_B3          TTTGGACTACTCGGGATCCGCCATCGATGAGATTAACAAGGATGTAATCCAACTCTCTGC
P20_J5          TTTGGACTACTCGGGATCCGCCATCGATGAGATTAACAAGGATGTAATCCAACTCTCTGC
                ******************************** ***************************

P20_J8          TGTCGCCACCTCTATCATCAAAATGCTGTTATGAAATATCTCATCTCATCATCATATCCT
P20_J1          TGTCGCCACCTCTATCATCAAAATGCTGTTATGAAATATCTCATCTCATCATCATATCCT
P20_B9          TGTCGCCACCTCTATCATCAAAATGCTGTTATGAAATATCTCATCTCATCATCATATCCT
P20_B2          TGTCGCCACCTCTATCATCAAAATGCTGTTATGAAATATCTCATCTCATCATCATATCCT
P20_J3          TGTCGCCACCTCTATCATCAAAATGCTGTTATGAAATATCTCATCTCATCATCATATCCT
P20_B1          TGTCGCCACCTCTATCATCAAAATGCTGTTATGAAATATCTCATCTCATCATCATATCCT
P20_J7          TGTCGCCACCTCTATCATCAAAATGCTGTTATGAAATATCTCATCTCATCATCATATCCT
P20_B3          TGTCGCCACCTCTATCATCAAAATGCTGTTATGAAATATCTCATCTCATCATCATATCCT
P20_J5          TGTCGCCACCTCTATCATCAAAATGCTGTTATGAAATATCTCATCTCATCATCATATCCT
                ************************************************************

P20_J8          CCATCCTCGAATCA
P20_J1          CCATCCTCGATCCA
P20_B9          CCATCCTCGATCCA
P20_B2          CCATCCTCGATCCA
P20_J3          CCATCCTCGATCCA
P20_B1          CCATCCTCGATCCA
P20_J7          CCATCCTCGATCCA
P20_B3          CCATCCTCGATCCA
P20_J5          CCATCCTCGATCCA
                **********  **
```

Key

Sequences highlighted in blue color represent position of Forward primer

Sequences highlighted in yellow color represent position of the reverse primer

ATG in green color represent the transcription start site/sequence

TGA in red color represent the transcription STOP site/sequence
