## Supplementary File S3 for "Diversity, Phylogenetic Relationships, And Expression Profiles Of Invertase Inhibitor Genes In Sweetpotato"

Figure S2:

Amino acid sequence alignment of invertase inhibitor proteins in sweetpotatoes. Sequences from other species were added as outgroups. **Accession numbers are preceded by 2 or 3-letters to identify the species. Iba, *Ipomoea batatas*; Itf, *Ipomoea trifida*; Itb, *Ipomoea triloba*; Sl, *Solanum lycopersicum*; Ath, *Arabidopsis thaliana; Csa, Camelina sativa.*** ***The four cysteine residues are conserved among all sequences and are numbered C1-C4. The PKF motif, conserved in dicots is underlined above PKF letters***

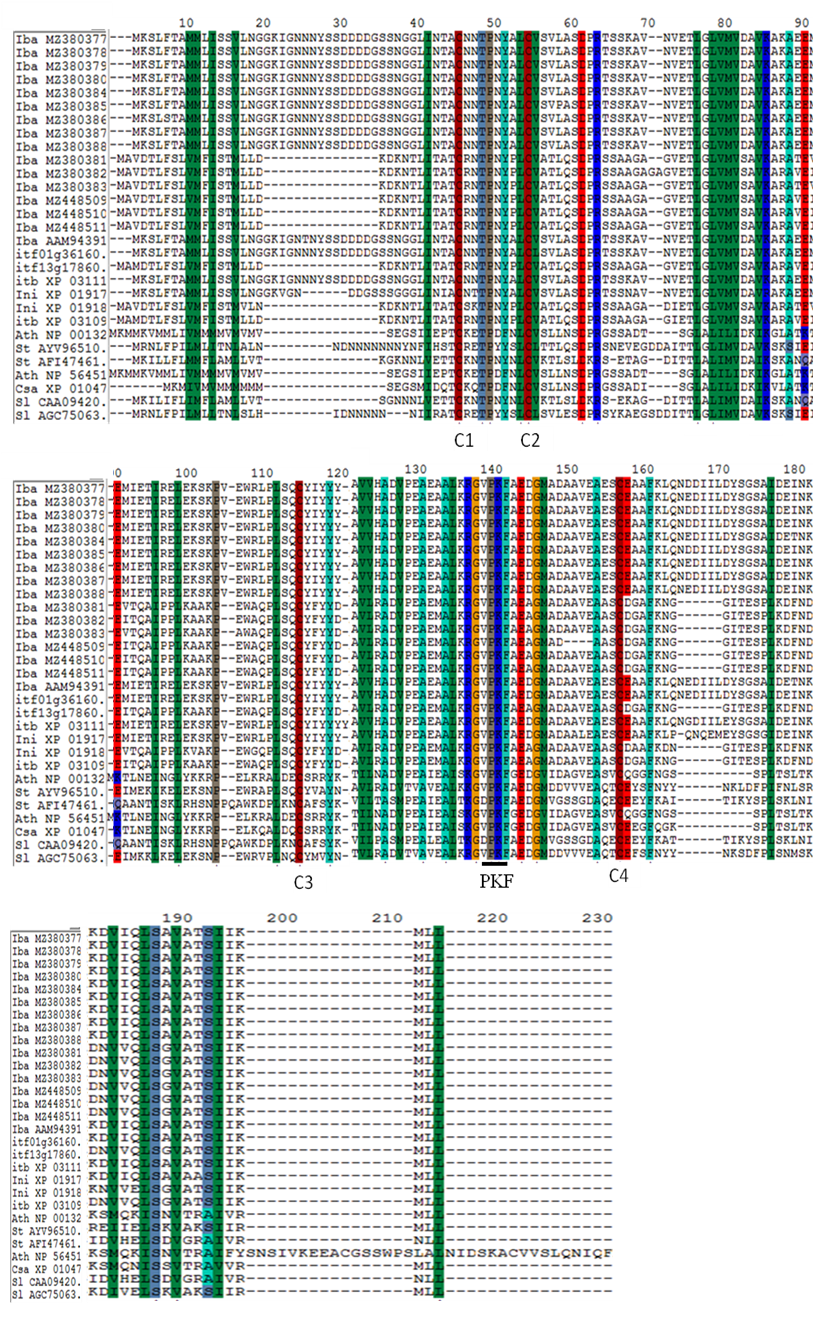

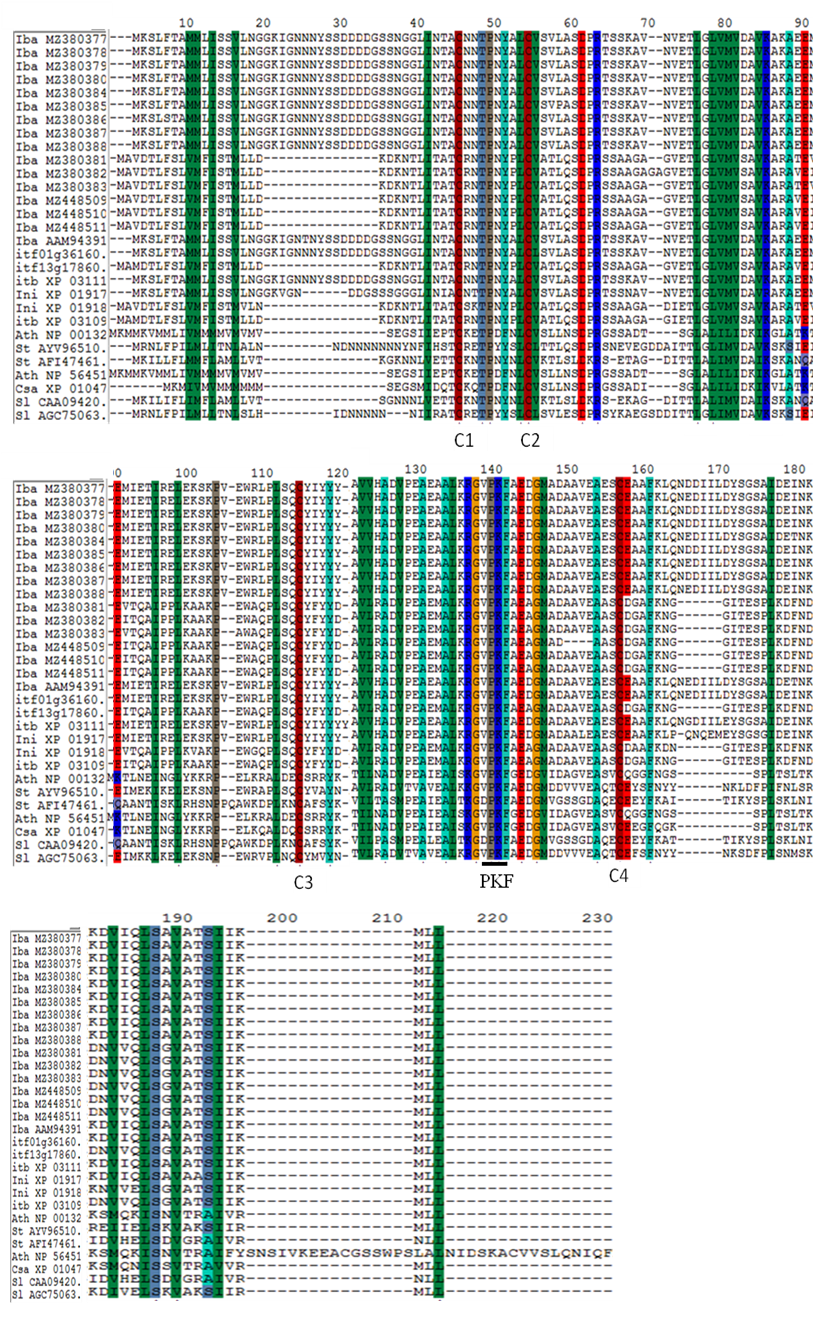

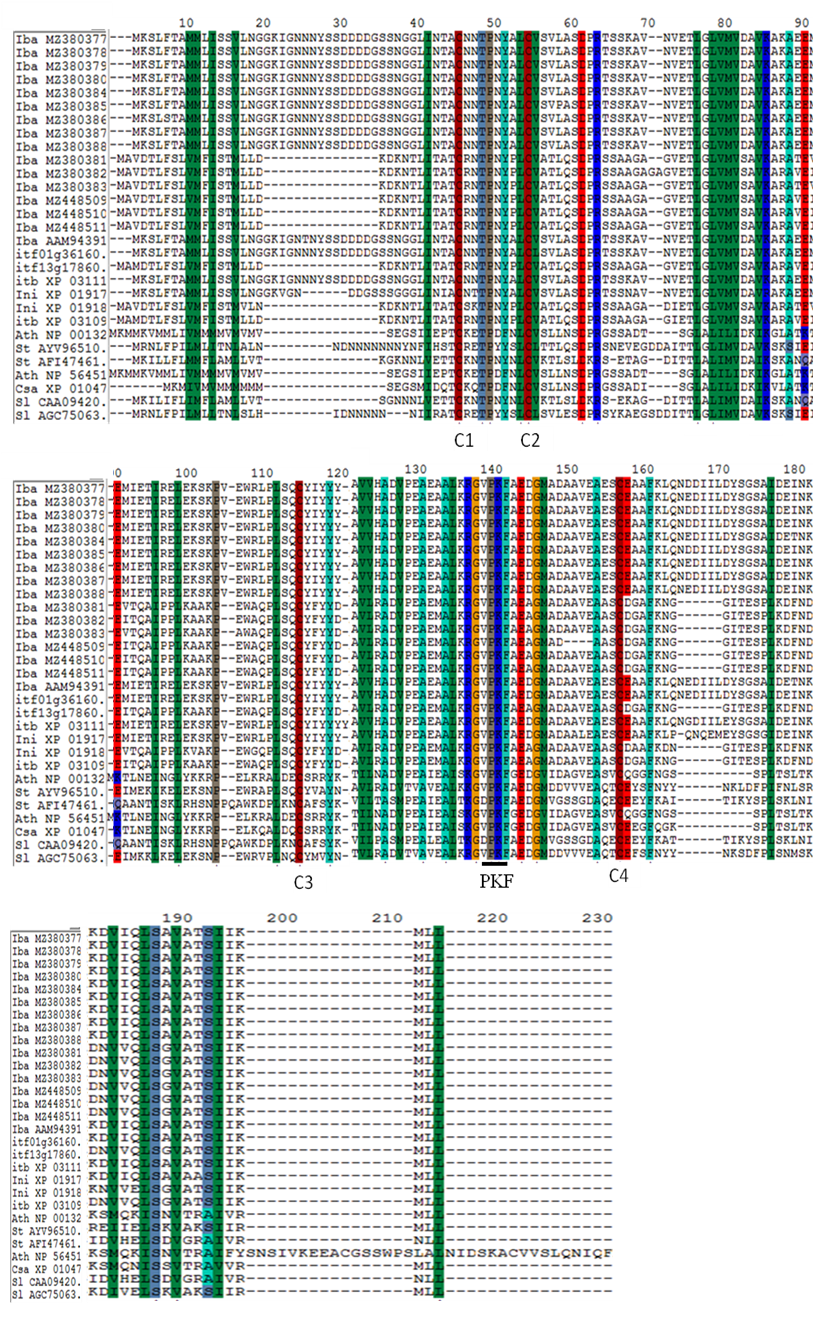
