## Supplementary Table S1 for "Diversity, Phylogenetic Relationships, And Expression Profiles Of Invertase Inhibitor Genes In Sweetpotato"

**Table S1: Properties of ITI proteins in sweetpotato**

| **Paralog** | **Source/variety** | **Accession No.** | **aa** | **Mw** | **pI** | **Homology (%)** | **E-value** |
| --- | --- | --- | --- | --- | --- | --- | --- |
| SPITI1 | Beauregard | MZ448509 | 168 | 17.88 | 5.13 | 54.21 | 9e-61 |
| SPITI1 | Beauregard | MZ448510 | 172 | 18.25 | 4.96 | 55.79 | 7e-65 |
| SPITI1 | Beauregard | MZ448511 | 172 | 18.25 | 4.96 | 55.79 | 7e-65 |
| SPITI1 | Jewel | MZ380381 | 172 | 18.24 | 4.96 | 55.79 | 4e-65 |
| SPITI1 | Jewel | MZ380382 | 174 | 18.38 | 4.69 | 55.21 | 9e-63 |
| SPITI1 | Jewel | MZ380383 | 172 | 18.18 | 5.13 | 55.26 | 3e-64 |
| SPITI2 | Tainong 57 | AAM94391.1 | 192 | 20.63 | 4.47 | 100 | 0.0 |
| SPITI2 | Beauregard | MZ380377 | 192 | 20.64 | 4.45 | 98.46 | 4e-143 |
| SPITI2 | Beauregard | MZ380378 | 192 | 20.65 | 4.47 | 98.46 | 4e-143 |
| SPITI2 | Beauregard | MZ380379 | 192 | 20.64 | 4.45 | 98.44 | 1e-42 |
| SPITI2 | Beauregard | MZ380380 | 192 | 20.64 | 4.45 | 98.44 | 1e-142 |
| SPITI2 | Jewel | MZ380384 | 192 | 20.64 | 4.47 | 99.48 | 4e-144 |
| SPITI2 | Jewel | MZ380385 | 192 | 20.62 | 4.45 | 97.92 | 3e-141 |
| SPITI2 | Jewel | MZ380386 | 192 | 20.59 | 4.47 | 98.44 | 1e-141 |
| SPITI2 | Jewel | MZ380387 | 192 | 20.64 | 4.45 | 98.44 | 1e-142 |
| SPITI2 | Jewel | MZ380388 | 192 | 20.65 | 4.47 | 98.96 | 4e-143 |
| ITI2 |  | XP_031112441 | 193 | 20.76 | 4.52 | 97.41 | 3e-139 |
| ITI2 |  | itf01g36160.t1 | 192 | 20.67 | 4.47 | 98.44 | 1e-142 |
|  |  | itf13g17860.t1 | 172 | 18.28 | 4.96 | 55.73 | 1e-65 |
|  |  | XP_019174712 | 185 | 19.77 | 4.57 | 87.05 | 3e-121 |
|  |  | XP_019180845 | 172 | 18.34 | 5.00 | 55.50 | 2e-64 |
|  |  | XP_031098734 | 172 | 18.29 | 4.96 | 55.21 | 3e-65 |
|  |  | NP_564516.2 | 205 | 22.33 | 8.44 | 42.98 | 1e-31 |
