## Supplementary Table S2 for "Diversity, Phylogenetic Relationships, And Expression Profiles Of Invertase Inhibitor Genes In Sweetpotato"

Predicted subcellular localization of ITI genes in sweetpotatoes

a. Predictions by TargetP

| **ID** | **Others** | **SP** | **MT** | **CH** | **TH** | **CS pos** | **pr** |
| --- | --- | --- | --- | --- | --- | --- | --- |
| MZ380380 | 0.056 | 0.9294 | 0.013 | 0.0003 | 0.0012 | 17-18, LNG-GK | 0.2427 |
| MZ380379 | 0.056 | 0.9294 | 0.013 | 0.0003 | 0.0012 | 17-18, LNG-GK | 0.2427 |
| MZ380378 | 0.0573 | 0.9278 | 0.0135 | 0.0003 | 0.0012 | 17-18. LNG-GK. | 0.2427 |
| MZ380388 | 0.0573 | 0.9278 | 0.0135 | 0.0003 | 0.0012 | 17-18. LNG-GK. | 0.2427 |
| MZ380387 | 0.056 | 0.9294 | 0.013 | 0.0003 | 0.0012 | 17-18. LNG-GK. | 0.2427 |
| MZ380386 | 0.3347 | 0.5868 | 0.0683 | 0.0041 | 0.0062 | 21-22. KIG-NN. | 0.2111 |
| MZ380385 | 0.041 | 0.9481 | 0.01 | 0.0003 | 0.0006 | 17-18. LNG-GK | 0.2425 |
| MZ380388 | 0.0057 | 0.9284 | 0.0131 | 0.0003 | 0.0012 | 17-18. LNG-GK | 0.2425 |
| MZ380377 | 0.056 | 0.9294 | 0.013 | 0.0003 | 0.0012 | 17-18. LNG-GK | 0.2425 |
| MZ448511 | 0.2725 | 0.7022 | 0.0006 | 0.0073 | 0.0173 | 28-29. ITA-TC | 0.3649 |
| MZ448510 | 0.2725 | 0.7022 | 0.0006 | 0.0073 | 0.0173 | 28-29, ITA-TC | 0.3649 |
| MZ448509 | 0.2533 | 0.7223 | 0.0006 | 0.0065 | 0.0174 | 28-29, ITA-TC | 0.3651 |
| MZ380381 | 0.2686 | 0.7084 | 0.0006 | 0.0096 | 0.0128 | 28-29ITA-TC | 0.3648 |
| MZ380383 | 0.2618 | 0.7202 | 0.0006 | 0.0067 | 0.0107 | 28-29, ITA-TC | 0.3652 |
| MZ380382 | 0.3106 | 0.663 | 0.0008 | 0.0095 | 0.0161 | 28-29, ITA-TC | 0.3656 |

- SP: for signal peptide,
- MT: for mitochondrial transit peptide (mTP),
- CH: for chloroplast transit peptide (cTP),
- TH: for thylakoidal lumen composite transit peptide (lTP),
- Other: for no targeting peptide (in this case, the length is given as 0).
- CS pos: for potential cleavage site
- Pr: for the probability associated with the predicted cleavage

b. Predictions by WoLF PSORT

| Accession No. /ID | Extr | Chl | Cyto | Mito | Golgi | Vacu | ER | Nuc | Plas |
| --- | --- | --- | --- | --- | --- | --- | --- | --- | --- |
| MZ380377 | 8 | 4 | 1 |  |  |  |  |  |  |
| MZ380378 | 8 | 4 | 1 |  |  |  |  |  |  |
| MZ380379 | 8 | 4 | 1 |  |  |  |  |  |  |
| MZ380380 | 8 | 4 | 1 |  |  |  |  |  |  |
| MZ380384 | 6 | 3 | 1 | 2 | 1 |  |  |  |  |
| MZ380385 | 7 | 4 | 2 |  |  |  |  |  |  |
| MZ380386 | 8 | 4 | 1 |  |  |  |  |  |  |
| MZ380387 | 8 | 4 | 1 |  |  |  |  |  |  |
| MZ380388 | 8 | 4 | 1 |  |  |  |  |  |  |
| MZ380381 | 1 | 8 | 2 | 1 |  |  |  |  |  |
| MZ380382 | - | 7 | 1 | 1 | - | 2 | 2 |  |  |
| MZ380383 | 1 | 8 | 1 | 1 | - | 2 |  |  |  |
| MZ448509 | 1 | 7 | 1 | 1 | - | - | 2 |  |  |
| MZ448510 | - | 6 | 1 | 1 | - | - | 2 |  |  |
| MZ448511 | - | 6 | 1 | 1 | - | 2 | 2 | 1 |  |
| AAM94391.1 | 7 | 4 | 1 | 1 | - | - | - | - |  |
| AYV96510.1 | 6 | 6 | - | - | - | - | 1.5 | - | - |
| AFI47461.1 | 5 | 1 | 1 | 1 | - | 1 | 3 | - | - |
| CAA09420.1 | 4 |  | 2 | 1 |  |  | 3 | 1 | 1 |
| AGC75063.1 | 3 | 8 |  |  |  | 2.5 | 2 |  |  |

c. Prediction by Phobius

| **Sequence ID** | **TM** | **SP** |  |
| --- | --- | --- | --- |
| MZ380377 | 0 | Y | N4-12c17/18o |
| MZ380378 | 0 | Y | N4-12c17/18o |
| MZ380379 | 0 | Y | N4-12c17/18o |
| MZ380380 | 0 | Y | N4-12c17/18o |
| MZ380384 | 0 | Y | N4-12c17/18o |
| MZ380385 | 0 | Y | N4-12c17/18o |
| MZ380386 | 0 | Y | N4-12c17/18o |
| MZ380387 | 0 | Y | N4-12c17/18o |
| MZ380388 | 0 | Y | N4-12c17/18o |
| MZ448509 | 0 | 0 | I |
| MZ448510 | 0 | 0 | I |
| MZ448511 | 0 | 0 | I |
| MZ380381 | 0 | 0 | I |
| MZ380382 | 0 | 0 | I |
| MZ380383 | 0 | 0 | I |

Visit <https://phobius.sbc.su.se/instructions.html> for interpretation of output
